# Differential fates of vertebrate Kazald gene quartet, from ancestral roles in skeletogenesis and regeneration to putative innovations in fish and birds

**DOI:** 10.1101/2025.06.21.660714

**Authors:** Sean D. Keeley, Rita Aires, Belfran Alcides Carbonell Medina, Claudia Marcela Arenas-Gómez, Alejandra Cristina López-Delgado, Jean Paul Delgado, Franziska Knopf, Shigehiro Kuraku, Tatiana Sandoval-Guzmán

**Author notes:** The author died prior to the submission of this paper (Jean Paul Delgado).

## Abstract

Salamanders are known for their incredible regenerative abilities, but translating findings to mammals is complicated by unannotated genes and unclear orthology. An example highlighting this difficulty was a discovery in the axolotl (*Ambystoma mexicanum*) of a regeneration-associated gene that has been identified as either *Kazald1* or *Kazald2*. Since orthology inference of genes across species is crucial to identifying gains and losses of gene functions, and thus if gene usage is likely to be consistent across species, we investigated the evolution of the axolotl genes using an extensive cross-species analysis. Molecular phylogeny inference conclusively identified the regeneration-associated gene as *Kazald2*, but also revealed an undescribed four-member Kazald gene family in jawed vertebrates. Moreover, synteny comparisons demonstrated that this family originated in the ancestral two-rounds of whole genome duplication. Additionally, we performed vertebrate-wide comparisons of Kazald gene expression profiles, employing available RNA-Seq which we validated in whole tissues of axolotl, zebrafish, and sharks. This uncovered seemingly ancestral connections conserved over jawed vertebrate evolution, such as *Kazald1* with skeletogenesis and odontogenesis and *Kazald2* with regeneration. It also suggested novel putative roles within specific lineages, including *Kazald3* in teleost fish skeletogenesis and *Kazald4* within avian brains. Our study thus demonstrates the establishment of a Kazald gene quartet in the jawed vertebrate ancestor, and elucidates the asymmetry of gene fates of its members, including deeply ancestral roles and comparably recent innovations. This provides a comprehensive report of this formerly undescribed gene family, offering a solid foundation for future studies of these genes in diverse species.

## Introduction

Salamanders have long been looked to as the golden standards of tetrapod regeneration due to their ability to completely regrow various structures, such as the limb, tail, brain, and eye, throughout their life (1). Meanwhile, such a high regenerative capacity is reduced in the majority of tetrapods, with even those that still possess this ability, such as frogs, losing it as they develop into adulthood (2, 3). However, many species outside of the tetrapod lineage possess a similar regenerative potential to salamanders, like the zebrafish (4), or even far greater abilities, such as the whole-body regeneration of hydra, planaria, and sea stars (5–7). This discrepancy in ability has driven much research into how regeneration proceeds in a variety of species in order to uncover the underlying mechanisms that drive this response. A common idea is that the ability to regenerate is an ancestral trait in vertebrates which has been lost or suppressed in most extant tetrapods (8). It may therefore be that the pathways used for regeneration in certain species are still present in regeneratively limited species like mammals, but are now utilized for different purposes and no longer activated by injury.

Several genes have been identified as promising candidates for reactivating these pathways and inducing successful regeneration, including *Prrx1*, *Msx2*, *Fetub*, and *Hdac1* (9–13). One particular gene, which was originally identified as *Kazal type serine peptidase inhibitor domain 1* (*Kazald1*) in the axolotl, is heavily expressed in the blastema of the regenerating limb but not in the developing limb bud, and the knockdown of it significantly slowed and reduced the regenerative response (9). This resulted in a great interest in this gene, as it appeared to be both specific to, and have a powerful effect on, regeneration. However, previous studies of regeneration in other species, such as the *Xenopus* frog and *Acomys* spiny mouse, do not report a similar upregulation of their *Kazald1* genes in the regenerating tissues (14, 15). Furthermore, different studies, and even literature reviews, in the axolotl were inconsistent in assigning a name to this gene, causing it to not always be labeled as *Kazald1* (11, 16, 17).

The latter inconsistency is of particular concern, as inaccurate gene identification and naming can create long-standing problems, especially for research focused on cross-species comparisons or on translating discoveries made in one species to humans. It could be that the same gene in various species, i.e., a set of orthologous genes, has been assigned different names across them. In this instance, any investigation of that gene in published research or online databases is greatly complicated, and previous discoveries related to it are prone to being overlooked. Alternatively, different genes in the compared species, i.e., paralogous or even non-homologous genes, may be given the same name. Examples of such confusion due the maintenance of different paralogs between species, known as “hidden paralogy”, can be seen in the Nodal and Wnt gene families (18, 19). When this occurs, any discoveries related to roles of a gene in one species may incorrectly be thought to likely be held by the paralogous or unrelated gene of the other species. Additionally, if the non-orthologous gene does not possess that function, then that role will be erroneously thought to have been either lost in the latter species or a novel development of the former species. This determination of how novel a gene function is becomes especially useful for predicting how translatable a finding in one species is likely to be to another.

Finally, incorrect identification also greatly complicates phylogenetic reconstructions of the evolutionary history of specific genes, and potentially even of the species that possess them. However, updating gene names [for easier use in research] to match their orthology can be very difficult, especially if the field has grown accustomed to the old nomenclature. This may result in large amounts of time during which incorrect and/or conflicting terminology will be used when discussing these genes (20). Therefore, it is important to correctly identify genes as early as possible in order to avoid greater confusion later on.

While a previous study of ours did make use of online phylogenetic tools to support our choice of *Kazald2* as the name of this axolotl gene (16), a more focused and exhaustive phylogenetic analysis was still needed to definitively classify its identity and relationship to the *Kazald1* genes of other species. Additionally, if the axolotl gene is truly not orthologous to the mammalian *Kazald1*, then it was important to determine if the axolotl still maintains an actual *Kazald1* gene, as well as if the non-orthologous gene is possessed by other species. This would enable us to accurately determine when any roles associated to these genes arose, such as that of *Kazald1* during skeletogenesis and odontogenesis in mammals (21) and that of the axolotl gene during regeneration (9). Similarly, it would uncover if these processes in different species have either evolved to not need any of these genes, or if non-orthologous genes have replaced them. However, untangling the phylogenetic relationships and characterizing ancestral roles required an in-depth analysis spanning distantly related species spread throughout the major vertebrate lineages.

In this work, we identified putative Kazald genes in approximately 60 vertebrate species spanning the tree-of-life, as well as several invertebrate deuterostome and protostome species representing major lineages within them. Molecular phylogeny analysis of these genes conclusively demonstrated that the regeneration-associated axolotl gene is in fact *Kazald2*. Even more importantly, it discovered the existence of a previously unknown gene family consisting of four members, including one gene, *Kazald4*, that has never been formally described. Synteny analysis uncovered that this gene family arose in the two-round whole genome duplication (2R-WGD) event that is ancestral to jawed vertebrates. We also provide a thorough account of which lineages still maintain specific Kazald genes, and by analyzing RNA-Seq expression data and performing RNA *in situ* hybridization and RT-qPCR, we have found tissues, including the brain and bones, and biological processes, such as regeneration and development, in which these genes are expressed across species. This will provide the research community with a fundamental knowledge of these genes, which would otherwise often be overlooked due to the large difficulties incurred by a lack of annotation and prior existing information. In this way, this work will also act as a solid foundation for future studies focused on individual Kazald genes and the mechanisms through which they work.

## Results

### Identification of Kazald genes in the axolotl

To determine if the regeneration-associated axolotl Kazald gene was orthologous to *Kazald1*, *Kazald2*, or a salamander unique gene, we searched through the axolotl transcriptome and genome via Reciprocal Best Hit (RBH) BLAST (22) using mouse *Kazald1*, and zebrafish *kazald2* and *kazald3* (**Table 1, Supp. Table 1**). This discovered four distinct genes spread across four different chromosomes. Additionally, the regeneration-associated axolotl gene (XM_069632371.1) was not the most similar one to mouse *Kazald1*, strengthening the hypothesis that it is not the *Kazald1* ortholog. However, the comparably low similarity of XM_069632371.1 to the zebrafish *kazald2* did not allow any obvious conclusions about potential orthology between these two genes. Furthermore, there being more Kazald genes in the axolotl than in the mouse and zebrafish combined greatly complicated the ability to identify evolutionary relationships with these limited species. Thus, we expanded our examination to include diverse species across the animal tree-of-life.

**Table 1.**
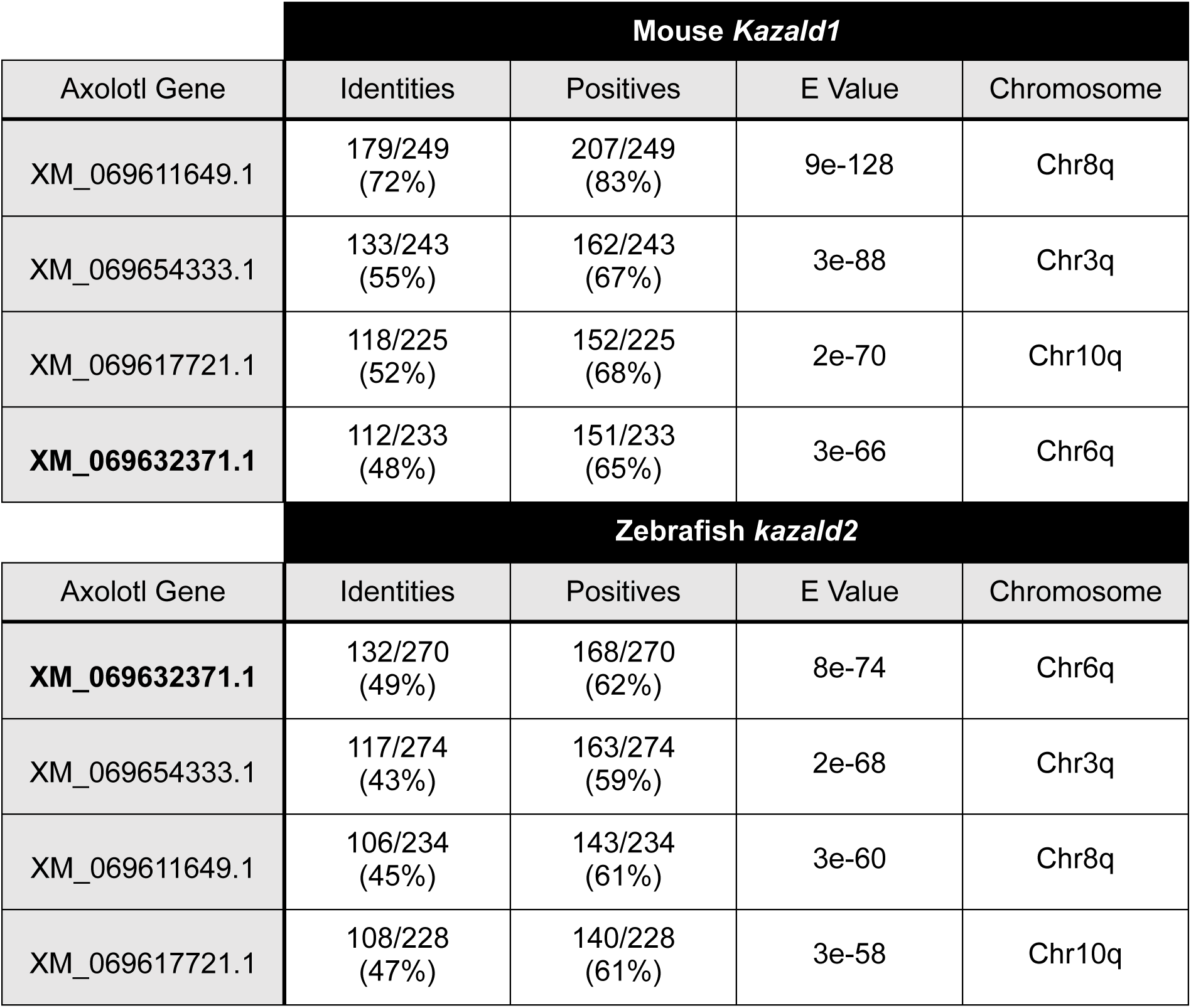
TBLASTN search reveals the presence of four axolotl genes with a high similarity to known Kazald genes. Identities are number of identical amino acids. Positives are aligned amino acids that are either identical or have similar chemical properties. E Value is the number of expected hits of similar quality that could be found just by chance. Chromosome indicates where the axolotl gene is located. Axolotl gene IDs taken from UKY_AmexF1_1 genome assembly (GCF_040938575.1). Regeneration-associated axolotl gene is bolded.

### The jawed vertebrate ancestor possessed four unique Kazald genes

This search particularly focused within jawed vertebrates, encompassing many species from its three major lineages: lobe-finned fish (sarcopterygians), ray-finned fish (actinopterygians), and cartilaginous fish (chondrichthyans). However, to completely explore the evolutionary history of these genes, representative species from the jawless fish lineage (cyclostomes), and several different invertebrate deuterostomes, including chordates and echinoderms, and protostomes, such as arthropods, mollusks, and annelids were included (**Supp. Table 2**). This analysis found Kazald genes in deuterostomes and protostomes, but not in cnidarians, placozoans, or sponges. Thus, the ancestral Kazald gene appears to have originated in bilaterians.

This analysis also revealed that most vertebrate lineages possessed multiple Kazald genes, with notable exceptions being cyclostomes, snakes, and placental mammals, which only had one (**Supp. Table 2**). However, few were comparable to the axolotl, although the sterlet and paddlefish contained even greater numbers. In contrast, most invertebrates only possessed one Kazald gene, except for some species from lineages known to have experienced whole genome duplications, like horseshoe crabs (23). This raised the idea that vertebrate Kazald genes were also established via genome duplication, specifically the 2R-WGD event ancestral to jawed vertebrates (gnathostomes) (24, 25). However, the lack of multiple Kazald genes in cyclostomes, which also experienced multiple genome duplications (26, 27), and no chondrichthyans having more than two Kazald genes, left open the possibility that at least some of these genes were unique developments within bony vertebrates (osteichthyans). To test this possibility, we performed a comprehensive phylogenetic analysis via maximum likelihood (ML) and Bayesian inference (BI) using the amino acid sequences of all our identified Kazald genes. This discovered that all jawed vertebrate Kazald genes were part of a single gene family, comprising four distinct clades (**Fig. 1**). This was supported by both phylogenetic methods, with their generated trees being extremely similar despite low posterior probabilities (PP) at basal nodes in the BI tree (e.g., 0.10 PP vs. 95 Bootstrap for the invertebrate Kazald node, and 0.08 PP vs. 62 Bootstrap for the clade grouping *Kazald1* with *Kazald3*). As BI with BAli-Phy is notoriously slow when handling large quantities of sequences (28), a potential explanation for these low values is that convergence at these basal nodes had not been reached when the analysis was terminated. This could be especially possible if genes with low placement support, such as those of the lamprey and hagfish, were varying in position across the set of trees due to ambiguity or conflict in their phylogenetic signal, making them rogue taxa (29, 30). As removal of rogue taxa can greatly increase resolution and branch support values within the consensus tree (31, 32), phylogenetic analyses were rerun using a subset of Kazald genes. This resulted in even higher support in both ML and BI trees for the four jawed vertebrate Kazald clades predicted by our initial large-scale analysis (**Supp. Fig. 1**).

**Figure 1.**
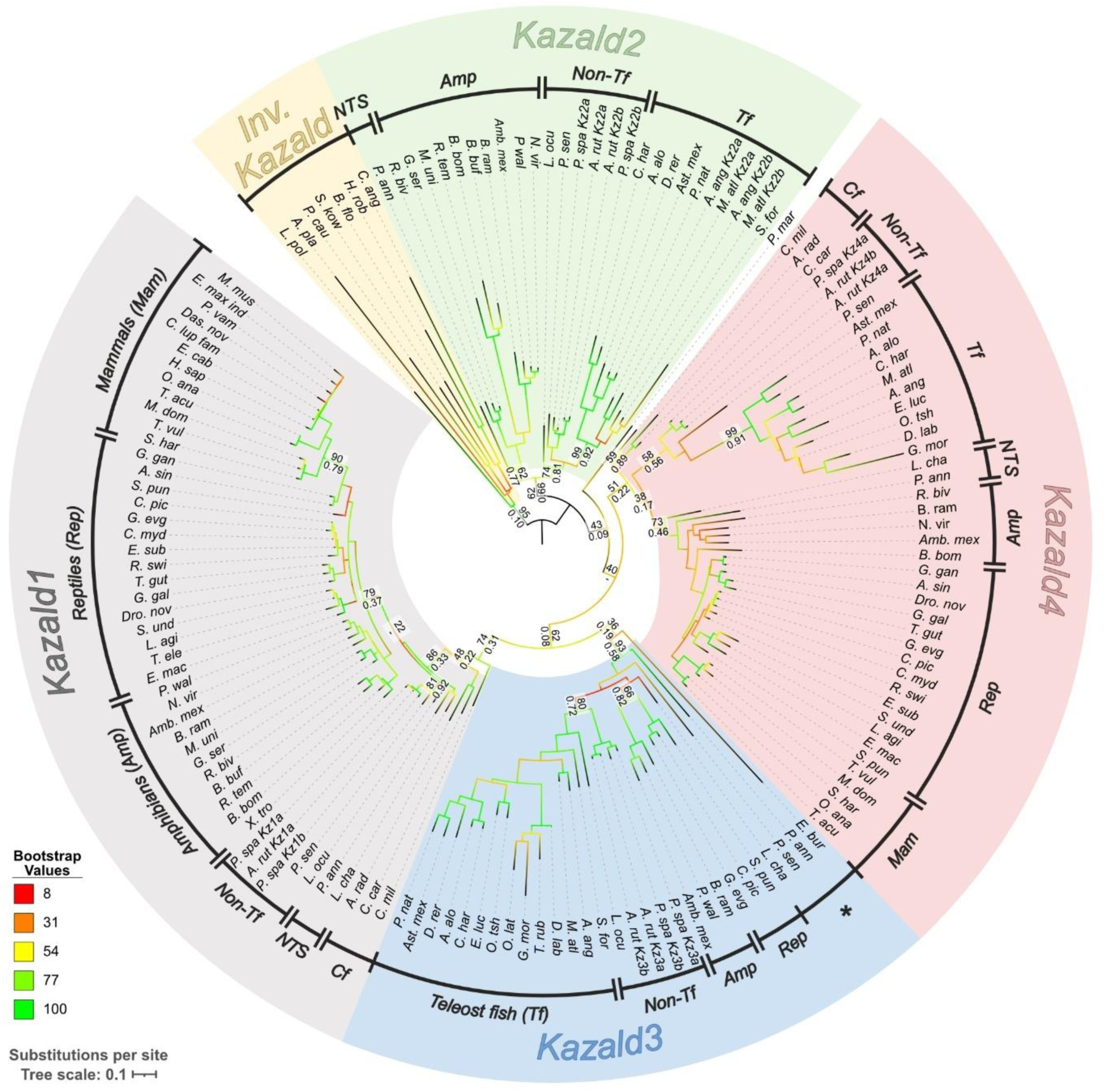
Phylogenetic tree of Kazald genes reveals that all vertebrate genes are split into four distinct clades. Displayed consensus tree was generated via RAxML using amino acid sequences. Branch colors indicate the bootstrap values at each node. Support values are shown at several key evolutionary nodes, e.g., at the base of mammals, amniotes, teleost fish, etc., or when the values between Maximum Likelihood (ML) and Bayesian Inference (BI) trees differ considerably. Layout of the support values are: ML bootstrap support via RAxML (top), BI posterior probabilities via BAli-Phy (bottom). Species are grouped together into major categories within each Kazald gene clade. Inv. = invertebrates, NTS = non-tetrapod sarcopterygians, Cf = cartilaginous fish, Non-Tf = non-teleost ray-finned fish. * marks a polyphyletic grouping containing a jawless vertebrate, two non-tetrapod sarcopterygians, and one non-teleost ray-finned fish. Abbreviated species names are listed in **Supplementary Table 2**. Kazald a and b versions created by whole genome duplications that occurred after the 2R-WGD event in certain fish lineages are marked next to the abbreviated species name. Scale bar corresponds to mean number of amino acid substitutions per site.

Three of these clades contained at least one previously validated gene from a major model species: *Kazald1* of human and mouse, and *kazald2* and *kazald3* of zebrafish. Therefore, the clades containing these genes were assigned the corresponding ID. Finally, to maintain consistency with the established sequential naming convention, we classified the remaining unlabeled clade as *Kazald4*, which to our knowledge has never been previously described. Importantly, the two chondrichthyan Kazald genes are *Kazald1* and *Kazald4*. As *Kazald2* is the outgroup to both these genes, but is possessed by several osteichthyan lineages, it was most likely present in the jawed vertebrate ancestor and later lost in chondrichthyans. Furthermore, the Kazald genes of lamprey and hagfish, representing the two major cyclostome lineages, were also part of different Kazald clades, indicating these genes are likely not orthologous, and existed prior to the cyclostome-gnathostome split. Taken together, the data supports the four jawed vertebrate Kazald genes most likely originating in the 2R-WGD event.

### Kazald gene quartet was established via two rounds of whole genome duplication

If the 2R-WGD event did generate the four gnathostome Kazald genes from a singular invertebrate Kazald gene ancestor, then the surrounding genes would have also been replicated. Furthermore, phylogenetic relationships between the resulting paralogs of each of these surrounding genes should be similar to that of the Kazald genes. Thus, we checked for conserved intragenomic synteny, i.e., the conservation of a similar order of paralogous genes (33, 34), between the genomic regions containing Kazald paralogs within a species.

We began by selecting a set of species that each contained as many of the four Kazald genes as possible, while also representing sarcopterygians, actinopterygians, and chondrichthyans. Teleost fish were excluded due to complications caused by their additional whole genome duplication. Ultimately, a species of salamander (*Ambystoma mexicanum*), turtle (*Mauremys mutica*), lungfish (*Protopterus annectens*), bichir (*Polypterus senegalus*), gar (*Lepisosteus oculatus*), and skate (*Amblyraja radiata*) were used, with four large genomic regions identified in each species. These were categorized as Syntenic Blocks A – D based on their contained Kazald and fibroblast growth factor subfamily D (FgfD) (35) gene. Regions of these blocks immediately surrounding the Kazald genes in a subset of these species are illustrated in **Fig. 2A**.

**Figure 2.**
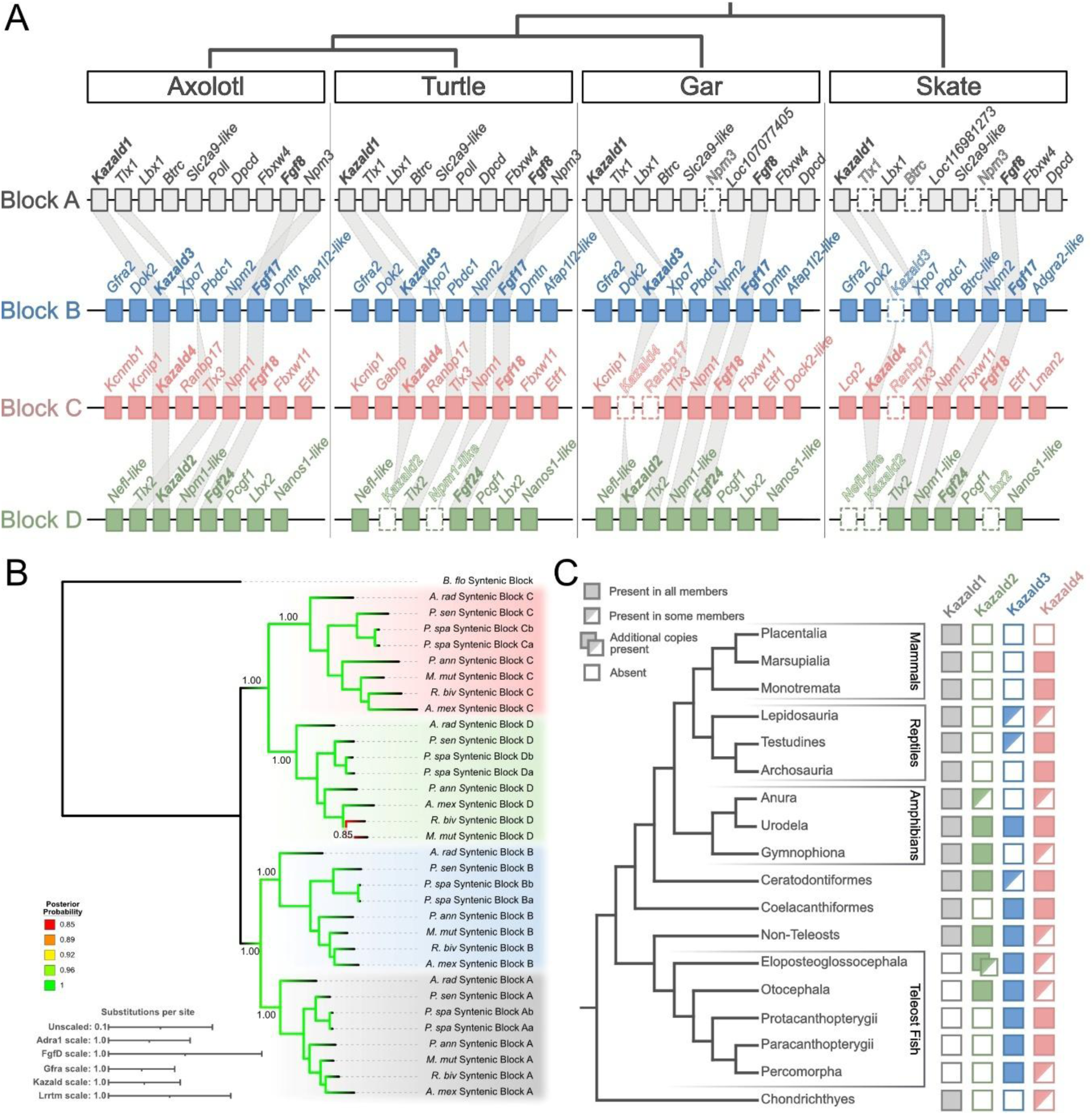
Conserved intra-/intergenomic synteny analysis supports the origin of the Kazald gene family in the 2R-WGD event ancestral to jawed vertebrates. A) Illustration of the syntenic blocks containing the Kazald genes in four representative species. Various paralogous genes are connected via gray lines. Genes lost in a species are drawn with a dotted outline and no internal color. Kazald genes are bolded. Fgf genes were used as the syntenic block anchor when a Kazald gene was absent, and thus are also bolded. B) Consensus tree generated via BAli-Phy partitioned analysis using the amino acid sequences of the five listed gene families (see also **Supp. Fig. 2**). Branch colors indicate the BI posterior probability at each node. Support values shown at nodes are BI posterior probabilities. Scale bar corresponds to mean number of amino acid substitutions per site for each of the used gene families. C) Simplified diagram of individual Kazald gene maintenance across vertebrates, along with the relationships of the different lineages. Colors of Kazald genes and the Syntenic Blocks correspond to each other. Dual Boxes for *Kazald2* in Eloposteoglossocephala represents the presence of *Kazald2a* and *Kazald2b* in the contained superorder Elopomorpha. Urodela is listed with the four Kazald genes present, as every examined species with a sequenced genome was found to possess all four genes.

This confirmed the existence of intragenomic synteny, with several families of known paralogous genes distributed across these syntenic blocks in each species. Furthermore, some of these paralogous genes, such as *Fgf8*/*17*/*18*/*24* and *T cell leukemia homeobox 1/2/3* (*Tlx1*/*2*/*3*), have previously been hypothesized to originate in the 2R-WGD event (36–38). Finally, the large-scale regions of the human and gar genome containing these syntenic blocks roughly correspond to Chordate Linkage Group I (CLGI) (39), further bolstering the idea that these regions are the products of the 2R-WGD event.

Apart from conserved intragenomic synteny within individual species, we observed conserved intergenomic synteny of the syntenic blocks across species. For example, the gene identified as *Kazald1* in a species was always associated with specific members of the aforementioned gene families, such as *ladybird homeobox 1* (*Lbx1*) and *Fgf8*, as well as with genes that lacked paralogs elsewhere in the genome, like *deleted in primary ciliary dyskinesia homolog (mouse)* (*Dpcd*). This conserved intergenomic synteny greatly supports there being exactly four Kazald clades in jawed vertebrates, as predicted in our initial tree (**Fig. 1**), and that genes within individual species were attributed to the appropriate clade. It also highlights other interesting evolutionary findings, such as *Fgf24*, which was thought to be lost in tetrapods, is still present in axolotl and turtle (40).

Finally, we analyzed the phylogenetic relationships of five gene families – adrenoceptor alpha 1 (Adra1), FgfD, GDNF family receptor alpha (Gfra), Kazald, and leucine rich repeat transmembrane neuronal (Lrrtm) – that still had a paralog present within each syntenic block in the majority of our selected species. While the exact structure of the gene trees sometimes differed, such as the double-fork of the Fgf tree vs. the stepwise splits of the Lrrtm tree, there was no discordance of which syntenic blocks were most closely related, e.g., Syntenic Block C was never more closely related to Syntenic Block A than it was to Syntenic Block D (**Supp. Fig. 2**). Additionally, no genes from a particular syntenic block were split apart, or mixed with genes of another syntenic block.

Thus, with no conflict between individual gene trees, we conducted a partitioned analysis using all of these gene families. This approach can more accurately reflect their true evolutionary histories by uncovering hidden support for internal relationships otherwise lost when only using individual gene families (41, 42). Additionally, using multiple genes acts as a better proxy for the complete syntenic blocks, and thus more accurately reflects the evolution of these genomic regions. Our partitioned multigene tree maintained the pairing of *Kazald1* (Syntenic Block A) with *Kazald3* (Syntenic Block B), now with even higher confidence (**Fig. 2B**). However, *Kazald4* (Syntenic Block C) and *Kazald2* (Syntenic Block D) now grouped together with very high confidence, matching what would be expected of four genomic regions originating from two sequential genome duplications. Thus, we can conclude that the Kazald gene family was created through the 2R-WGD event, and therefore all four genes were present in the gnathostome ancestor.

### Kazald4 is highly expressed in avian brains

With the phylogeny of the syntenic blocks predicted with high confidence, an overview of Kazald gene maintenance across vertebrates could be achieved, revealing that the conservation of Kazald genes differs greatly across lineages (**Fig. 2C**). However, some Kazald genes were more likely to be maintained than others, with *Kazald1* and *Kazald4* being the first and second-most consistently conserved genes across tetrapods, respectively. Surprisingly, despite *Kazald4* being present in salamanders, lizards, birds, and even non-placental mammals, we could not find a formal report of it or its expression. This was unexpected for such a widely conserved gene, and so we examined public gene expression atlases for potential expression profiles (43). This found that *KAZALD4* (current annotation: *LOC769726*) is highly expressed in many parts of chicken brain. We investigated if this expression was conserved in other avian species by analyzing published RNA-Seq data. This found that *Kazald4* was similarly highly expressed in the whole brain of emu, and various parts of the brain of zebra finch (**Fig. 3A**). As these species span the entire avian lineage, *Kazald4* is most likely expressed in the brains of birds in general (**Fig. 3A’**). However, *Kazald4* was not similarly expressed in data generated from brains of other bony vertebrates (**Fig. 3B**). This included alligators of the order Crocodilia, the most closely related extant lineage to birds, indicating that its expression in avian brains is likely a unique adaptation in this lineage. However, we could not disregard the possibility that this adaptation arose earlier within the archosaur lineage, but was then lost in the branch leading to Crocodilia.

**Figure 3.**
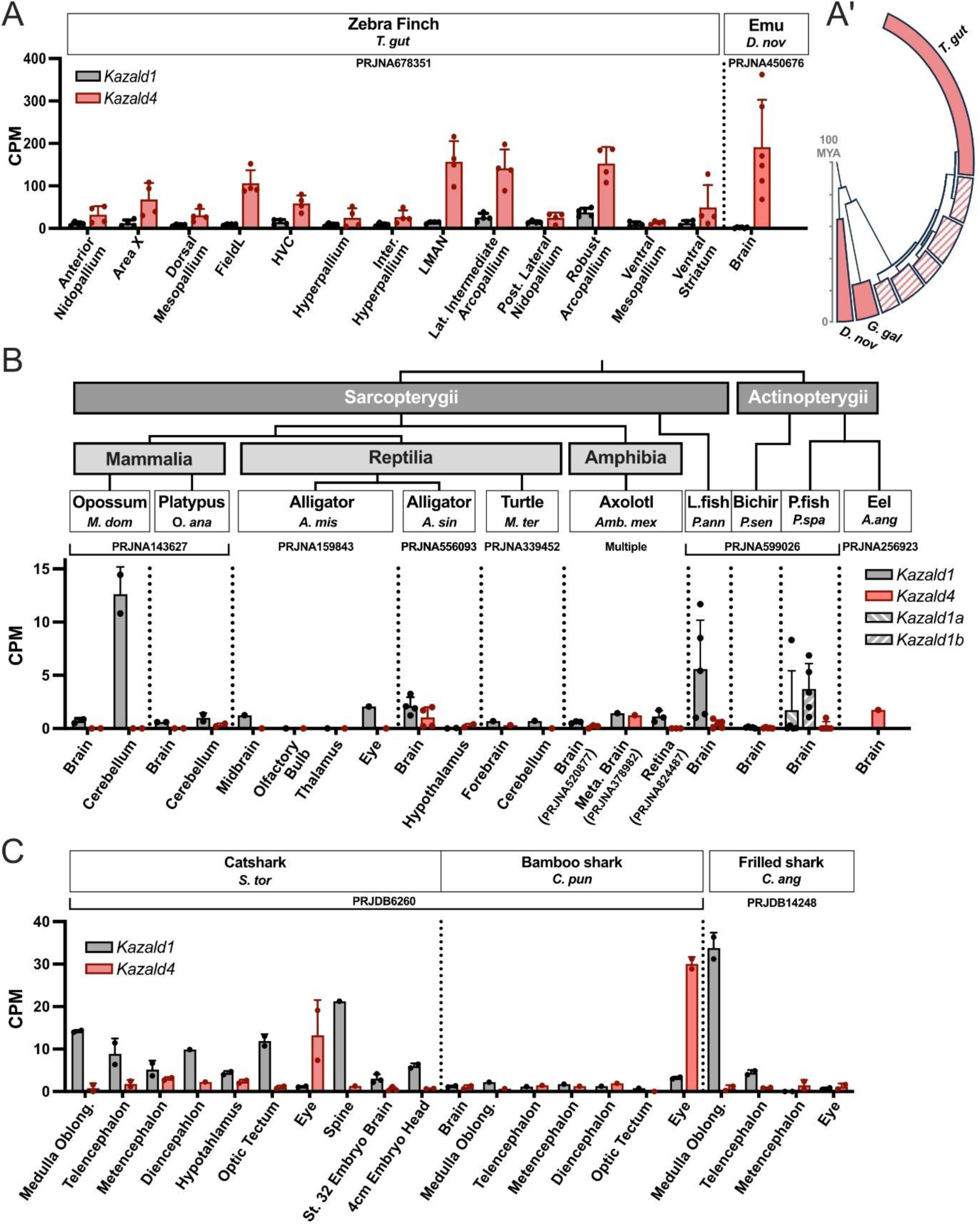
*Kazald4* is highly expressed in avian brains, but not in the brains of other species. A) Quantification of Kazald gene expression in different parts of the zebra finch brain and in whole emu brain. A’ shows a simplified evolutionary tree of birds (based on (45)); clades containing species shown to express *Kazald4* in the brain are completely filled in, while remaining clades have a striped pattern. B) Quantification of *Kazald1* and *Kazald4* gene expression in the brains and retinas of non-avian osteichthyans. A simplified overview of the evolutionary relationships of the examined species are displayed above the gene expression graph. *Kazald1* was duplicated in the lineage leading to paddlefish (P.fish) with no subsequent loss, creating a *Kazald1a* and *Kazald1b*. L.fish = Lungfish. C) Quantification of Kazald gene expression in different parts of the brain and associated neural tissues of several chondrichthyans. PRJ IDs indicate the publicly available RNA-Seq datasets which generated the raw data for the listed tissues. Dots represent biological replicates in examined datasets, error bars represent standard deviation when calculable. CPM = Counts Per Million.

Interestingly, data from some shark species demonstrated *Kazald4* expression in eyes, a region of which constitutes part of the central nervous system (44) (**Fig. 3C**). However, it could not be determined if *Kazald4* was specifically expressed in the neural tissue of the eye. Since our genetic and proteomic analysis of the four Kazald genes found strong similarities in their exon-intron structure, layout of protein domains, and the majority of their 3D protein structure (**Supp. Fig. 3 and Supp. Fig. 4**), we wondered if a different Kazald gene could be performing a similar role to it, thus explaining the lack of *Kazald4* expression in non-avian brains. Investigation of the other Kazald genes revealed noticeable levels of *Kazald1* in at least parts of the brain of a few jawed vertebrate species, including some sharks, lungfish, opossum, and zebra finch (**Fig. 3A-C**). However, most species did not express this gene, nor any other Kazald gene, in their brains (**Supp. Fig. 5**). Thus, it appears that, except for birds, there is no strong connection of any Kazald gene to the brain that is consistent within a wider lineage.

This lack of consistent *Kazald4* expression patterns ultimately extended across non-avian lineages. Some individual species of lungfish and sharks expressed this gene in various tissues, but these expression profiles were not recapitulated within other species within their lineage or closely related ones (**Supp. Fig. 6A and B)**. Furthermore, most species did not express much, if any, *Kazald4* in their examined tissues, nor during major biological processes like development or regeneration (**Supp. Fig. 6C and D)**. Thus, while *Kazald4* is the second-most common Kazald gene across jawed vertebrate lineages, it appears to lack a conserved expression profile that could explain this continued preservation.

### Kazald1 expression has ancestral ties to the skeleton and teeth

Previous investigations have associated *Kazald1* to skeletal development, finding expression in maturing osteoblasts and odontoblasts during bone and teeth mineralization in the mouse (21). However, due to the differential expression between lineages found in our analysis of *Kazald4,* we investigated if this role in skeleton and tooth development may be deeply ancestral, or if it was a later innovation in the lineage leading to mammals.

Published RNA-Seq reads of skeletal and dental tissues from a variety of species of diverse lineages were analyzed. Starting in mouse, we examined data from developing forelimbs of embryos and regenerating digits of adults, as both undergo periods of rapid ossification (46, 47). The former displayed a clear increase in *Kazald1* expression over limb development (**Fig. 4A**), importantly reflecting RNA *in situ* hybridization experiments from published studies (21). Meanwhile, the latter uncovered a peak of *Kazald1* expression in regenerating digit tips (R) at 14 days post amputation (dpa), which was not as striking in non-regenerating digits (NR), and is thus likely corresponding to the period of greatest bone reformation during regeneration (46). This reinforces the link of *Kazald1* with skeletal ossification, and extends it to adult mouse tissues.

**Figure 4.**
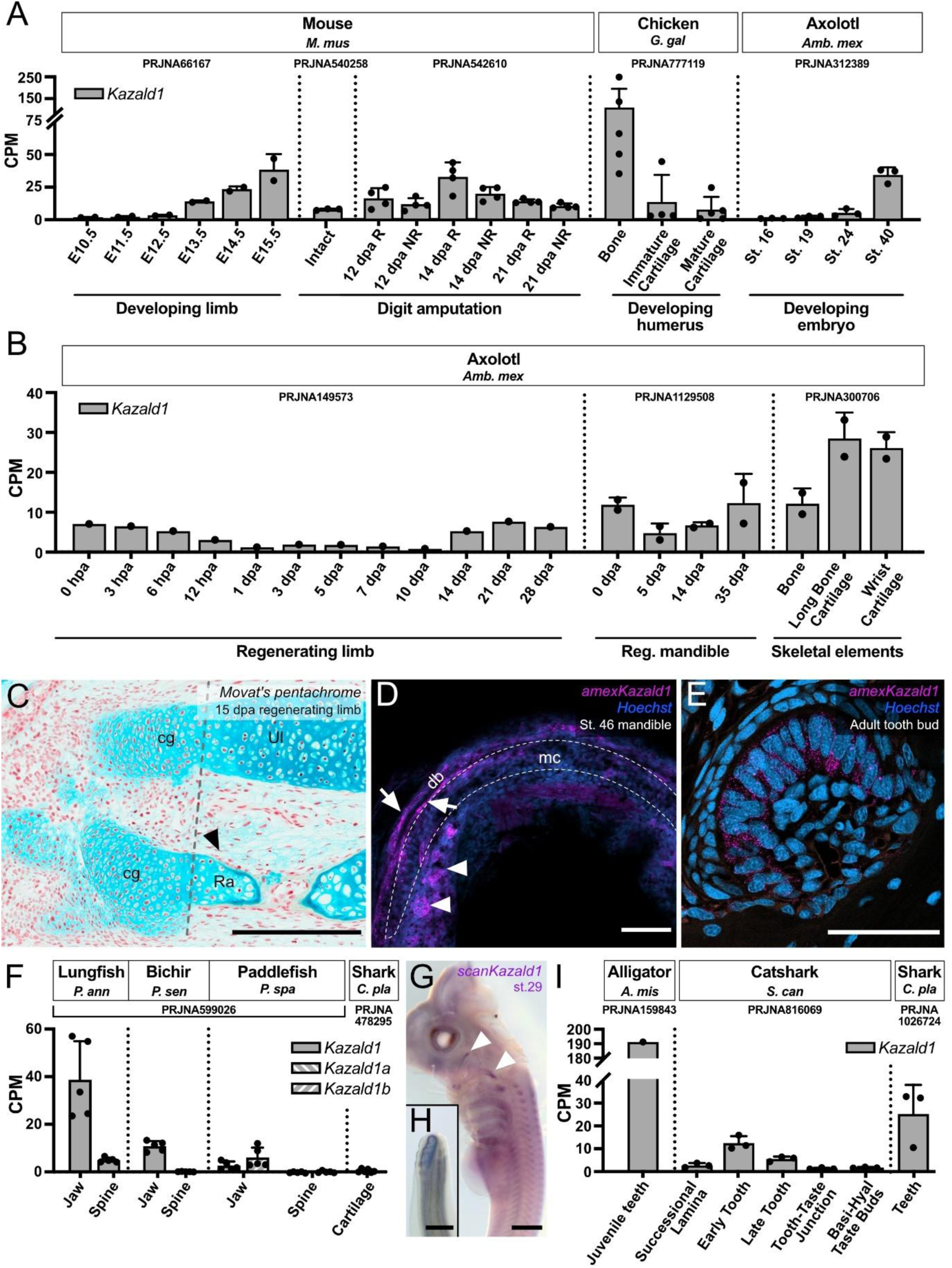
*Kazald1* is expressed during skeletal and tooth development in tetrapods, and potentially in all jawed vertebrates. A) Quantification of *Kazald1* expression in the developing whole limbs of embryonic mice, the intact, regenerating (R), and non-regenerating (NR) digit tips of amputated adult mice, the developing humerus of stage HH36 embryonic chickens, and whole embryos of different developmental stages of the axolotl. B) Quantification of *Kazald1* expression in adult axolotl during limb and mandible regeneration, and in the intact bone and cartilage of limbs. C) Movat’s pentachrome staining of 15 dpa juvenile axolotl limb. Scale bar: 500 μm. D) Z average projection of whole mount HCR for *Kazald1* in the lower jaw of stage 46 larval axolotl. Dashed line indicates location of Meckel’s cartilage (mc). Arrows indicate *Kazald1* expression associated to the ossifying dentary bone (db). Arrowheads indicate *Kazald1* expression associated to a tooth bud. Scale bar: 100 μm. E) HCR of *Kazald1* in a sectioned tooth bud of adult axolotl. Scale bar: 100 μm. F) Quantification of *Kazald1* expression in the jaw and spine of the lungfish, bichir, and paddlefish, and in the cartilage of bamboo shark. *Kazald1* was duplicated in the lineage leading to paddlefish with no subsequent loss, creating a *Kazald1a* and *Kazald1b*. G) Whole mount *in situ* for *Kazald1* in stage 29 catshark embryo head and upper trunk. Arrowheads indicate expression associated with the pharyngeal arches. Scale bar: 1 mm. H) Tail tip of embryo in G. Scale bar: 500 μm. I) Quantification of *Kazald1* expression in the tooth of juvenile alligator, at different stages of tooth development in adult catshark, and teeth of adult bamboo shark. Non-tooth tissue is included as controls for expression in other areas of the jaw of the catshark. PRJ IDs indicate the publicly available RNA-Seq datasets which generated the raw data for the listed tissues. Dots represent biological replicates in examined datasets, error bars represent standard deviation when calculable. CPM = Counts Per Million, St. = stage, dpa = days post amputation, hpa = hours post amputation.

We next examined *Kazald1* expression in RNA-Seq data from other sarcopterygians, and found a strong expression of *Kazald1* in bone of the developing humerus of embryonic chicken (**Fig. 4A**). Furthermore, this gene was expressed specifically in bone and not in cartilage, strongly suggesting that, similar to mouse, expression is linked to osteoblasts. Data from the axolotl also displayed an increase in *Kazald1* expression in the final stages of embryonic development, as well as during later stages of limb and jaw regeneration (**Fig. 4A and B**), both times in which extensive growth of skeletal elements occurs (16, 48).

However, in contrast to chicken, axolotl data revealed substantial *Kazald1* expression in both bony and cartilaginous regions of the intact limb. Expression in cartilage could explain why axolotl stage 40 whole embryos already express this gene even though ossification only starts occurring 10-12 days later in the jaw (49–51). Similarly, expression in cartilage could be why *Kazald1* returns to baseline levels in regenerating limbs by 15 dpa, despite the regenerated skeletal elements not being mineralized yet (**Fig. 4C**). Finally, axolotl limb chondrocytes were reported to express certain genes considered to be osteoblast-specific in amniotes (52), which might include *Kazald1*. Interestingly though, this may differ between parts of the skeleton, as our *in situ* hybridization chain reaction (HCR) of the embryonic jaw clearly shows expression in the ossifying dentary (db) bone rather than in the chondrocytes of the Meckel’s cartilage (mc) (**Fig. 4D, white arrows**).

Outside tetrapods, examination of RNA-Seq data from lungfish, a non-tetrapod sarcopterygian, and bichir and paddlefish, two non-teleost actinopterygians, revealed a lack of expression of *Kazald1* in their spines, but variable amounts in their jaws (**Fig. 4F**). Interestingly, their spines are largely cartilaginous, while their jaws are ossified (53–59). Therefore, the difference in expression between these skeletal elements suggests that, unlike the axolotl but similar to amniotes, *Kazald1* expression in these osteichthyan fish could be limited to ossified regions. However, the expression of *Kazald1* across several lungfish tissues (**Supp. Fig. 7A**), complicates a conclusive association to skeletogenesis in this species. Despite this, a connection to skeletogenesis specifically during development, even of cartilaginous elements, might still be ancestral to jawed vertebrates, as our whole-mount *in situ* hybridization (WISH) of catshark embryos seemed to show that *Kazald1* is associated with the pharyngeal arches that develop into the jaws (**Fig. 4G, white arrowheads**). However, it may instead coincide with certain cranial nerves (60), which would agree with its identified expression in the adult brain of another species of catshark.

The variable levels of *Kazald1* in these sarcopterygian and actinopterygian fish jaws could also indicate this gene being expressed in their teeth, the other major mouse tissue that expresses *Kazald1* (21). These fish differ greatly in dentition, with lungfish possessing large and continuously growing tooth plates (61), bichir exhibiting many conical teeth (62), and paddlefish lacking teeth as adults (63). Thus, the correlation of *Kazald1* expression with the extent of dentition raises the possibility that the connection to tooth development also exists in these early diverging species. We thus examined published RNA-Seq data of the teeth of several species.

This revealed a high expression in alligator teeth, some expression in the early developing tooth of the shark *Scyliorhinus canicula*, and variable expression in teeth of the shark *Chiloscyllium plagiosum* (**Fig. 4I**). Moreover, our RNA *in situ* hybridization study of *S. canicula* embryos found it expressed in developing caudal dermal denticles (**Fig. 4H**), which express many tooth-associated genes (64). Finally, our HCR staining of embryonic and adult axolotl jaws found robust expression within their teeth (**Fig. 4D, white arrowheads**, **and E**). Ultimately, these findings suggest an association between *Kazald1* and odontogenesis in tetrapod and chondrichthyan lineages, indicating this connection is likely ancestral to jawed vertebrates.

### Kazald3 is prone to loss, but has potentially replaced Kazald1 in teleost fish

In contrast to *Kazald1*, *Kazald3* is quite prone to loss, being completely absent in chondrichthyans and commonly lost in sarcopterygians (**Fig. 2C**). This absence in common tetrapod models makes it difficult to identify an ancestral expression profile, or determine if it even has one. RNA-Seq data from axolotl barely expressed this gene in any of a variety of tissues, which was also the case in data from the turtle *Malaclemys terrapin* and several non-tetrapod/teleost fish (**Supp. Fig. 7 and 8**). Expression in axolotl did slightly increase in the limb and jaw during regeneration, but a similar upregulation was not observed in data from regenerating brain or retina (**Fig. 5A**). To find what tissues expressed *Kazald3* during limb regeneration, we examined spatial transcriptomic data of regenerating axolotl limb previously published by our lab (65). This revealed sparse *Kazald3* expression predominantly spread across the limb dermis, which could explain the lack of upregulation in regenerating brain and retina.

**Figure 5.**
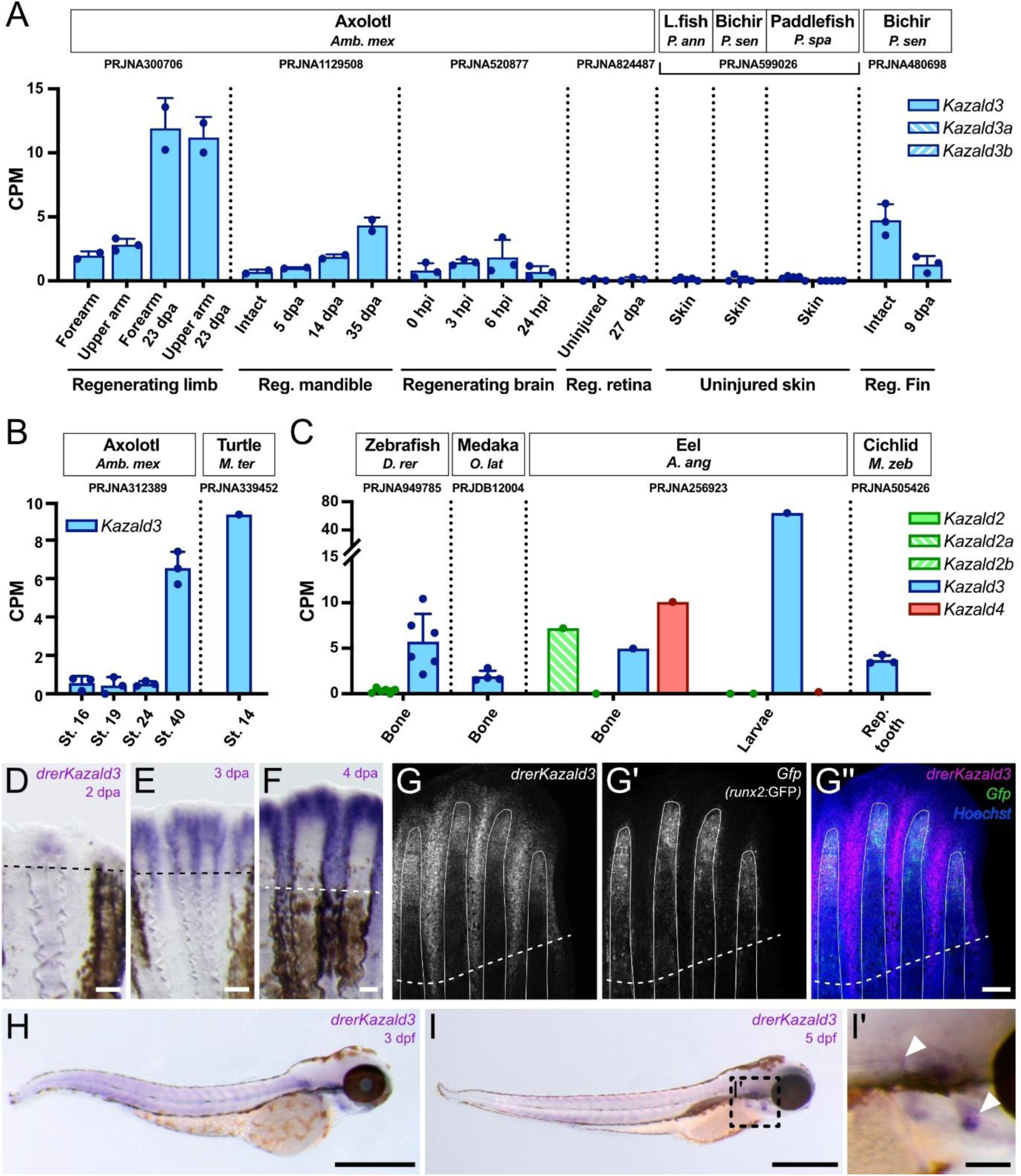
*Kazald3* may have a connection to skeletogenesis and regeneration in teleost fish, but has no clear role in other lineages. A) Quantification of *Kazald3* expression in regenerating limbs, lower jaw, brain, and retina of adult axolotl, the uninjured skin of non-tetrapod/teleost fish, and in regenerating fins of adult bichir. *Kazald3* was duplicated in the lineage leading to paddlefish with no subsequent loss, creating a *Kazald3a* and *Kazald3b*. L.fish = Lungfish B) Quantification of *Kazald3* expression in whole embryos of developing axolotl and turtle. C) Quantification of Kazald gene expression in the bone of adult zebrafish, bone of adult medaka, adult bone and the larvae of eel, and developing tooth of the mbuna cichlid. *Kazald2* was duplicated in the lineage leading to eel with no subsequent loss, creating a *Kazald2a* and *Kazald2b*. D – F) Whole mount *in situ* hybridization of *kazald3* in regenerating caudal fins of adult zebrafish from 2 to 4 dpa. Dashed line indicates the amputation planes. Scale bars: 100 μm. G – G’’) HCR of *kazald3* (G), *gfp* (G’), and their overlap (G’’) in 4 dpa regenerating caudal fin of adult *runx2*:GFP transgenic zebrafish. Dashed line indicates the amputation plane; solid white lines outline individual fin rays. Scale bar: 100 μm. H – I) Whole mount *in situ* hybridization of *kazald3* in 3 (H) and 5 (I) dpf zebrafish embryos. Scale bar: 500 μm. I’) Inset displaying the boxed region of the embryo in I. Arrowheads indicate areas of *kazald3* expression that associate to the locations of ossifying skeletal elements (i.e., the opercle and cleithrum). Scale bar: 100 μm. PRJ IDs indicate the publicly available RNA-Seq datasets which generated the raw data for the listed tissues. Dots represent biological replicates in examined datasets, error bars represent standard deviation when calculable. CPM = Counts Per Million, St. = stage, dpa = days post amputation, hpi = hours post injury, Reg. = regenerating, Rep. = replaced.

Meanwhile, *Kazald3* was only scarcely expressed at most in RNA-Seq data from uninjured skin of other species, which could imply that expression in skin only occurs during regeneration. However, analysis of data from bichir fin found a decrease in *Kazald3* expression during regeneration, and so an increase in expression during regeneration may instead be specific to axolotls/salamanders. Besides regeneration, *Kazald3* was also lowly expressed in certain embryonic stages of axolotl and turtle (**Fig. 5B**). Thus, a role for *Kazald3* during development may exist within the few tetrapod species still possessing it, but future study is needed to confirm this. Ultimately then, no strong or conserved links were detected between *Kazald3* and any tissue or biological process within the few sarcopterygian species that still maintain it, likely explaining why this gene is so prone to being lost in this lineage.

In a major departure from these other lineages though, *Kazald3* is the only consistently conserved Kazald gene in teleost fish (**Fig. 2C**). This is also associated with an extremely unusual loss of *Kazald1*, which is otherwise conserved in all other lineages. Due to the ancestral connection of *Kazald1* with skeletogenesis and odontogenesis, we hypothesized that *Kazald3* may have replaced *Kazald1* for these roles in teleosts.

Analysis of published RNA-Seq data from three distantly related teleost fish was inconclusive in linking *Kazald3* with bones (**Fig. 5C**). In support of our hypothesis, expression levels in intact teleost bones were similar to that of *Kazald1* in adult mouse and axolotl intact bones (**Fig. 4A and B**). Furthermore, *Kazald3* was heavily expressed in eel larvae, a stage with bone ossifying in the jaw, operculum, and caudal fin, and preceding extensive bone ossification throughout the body (66). However, *Kazald2a* and *Kazald4* being more highly expressed than *Kazald3* in the bone of the eel could indicate that other and/or multiple Kazald genes were instead associated to the bone in certain teleost fish lineages. This, combined with the overall low expression of *Kazald3* in the adult bones of each fish species, prevented making any conclusions regarding its role in teleost skeleton development.

We thus examined public single-cell zebrafish gene expression atlases, which revealed a clear expression of *kazald3* in osteoblasts, but also in non-osteoblast blastema cells and the basal layer of the wound epidermis in the regenerating fin at 3 dpa (67). Our WISH analysis of multiple days of fin regeneration found expression throughout the ray tissue and inter-ray areas of the regenerate, corroborating this single-cell data (**Fig. 5D-F**). Furthermore, HCR staining of 4 dpa fins found *kazald3* expression in the fin ray overlapping with the area of heavy osteoblast concentration, identified by *green fluorescent protein* (*gfp*) expression driven by a *RUNX family transcription factor 2* (*runx2*) promoter (68), supporting a connection with new bone ossification (**Fig. 5G**). To investigate if expression was limited to regenerating tissue, we examined developing zebrafish larvae. Our WISH analysis showed that *kazald3* also overlapped with the region of bone ossification in the developing head at 3- and 5-days post fertilization (dpf) (**Fig. 5H and I**), such as in the area of the operculum at 5 dpf (**Fig. 5I and I’**).

Thus, our work suggests that *kazald3* has replaced *Kazald1* during bone formation in zebrafish, and potentially across teleosts in general, explaining its uniquely strong conservation within this lineage. We also show that it likely possesses roles outside the skeleton in teleosts, as demonstrated by its expression in non-osteogenic tissue in regenerating fins.

### Kazald2 possesses an ancestral role in bony vertebrate regeneration

Our extensive phylogenetic analysis demonstrated that the regeneration-associated axolotl gene which prompted this investigation is *Kazald2*. However, it also uncovered widespread loss of *Kazald2* across lineages (**Fig. 2C**). Such common loss could indicate *Kazald2* lacked a useful role in these lineages, or was replaced by a different gene(s). Alternatively, its loss could be connected to the loss in regenerative potential of many of these lineages, although it is unclear which loss would have occurred first in this case. To investigate if *Kazald2* has an ancestral connection to regeneration, and its correlation to regenerative potential, we analyzed published RNA-Seq data from regenerating tissues of distantly related species that still maintain *Kazald2*.

In axolotl, a time course of limb regeneration demonstrated a large increase in expression from negligible levels in the intact limb up to a high peak in the early bud blastema stage at 7 dpa (**Fig. 6A**). This subsequently decreased back to basal levels, which agrees with published RNA *in situ* expression patterns (9). Examination of data from regenerating fins and tails of non-tetrapod sarcopterygian (two species of lungfish) and non-teleost actinopterygian (one species of bichir) fish revealed these species also greatly upregulate *Kazald2* from intact levels (**Fig. 6B**). This strongly supports a connection of *Kazald2* to regeneration since at least the last common ancestor of all bony vertebrates. Additionally, data from salamanders found that *Kazald2* is highly expressed during regeneration in many bony and soft tissues, including the mandible, spine, and retina (**Fig. 6C**). Expression in the brain was variable, but published HCR staining of *Kazald2*, although referenced as *Kazald1*, has demonstrated weak to strong expression in axolotl regenerating brain from 2 to 7 dpi (69).

**Figure 6.**
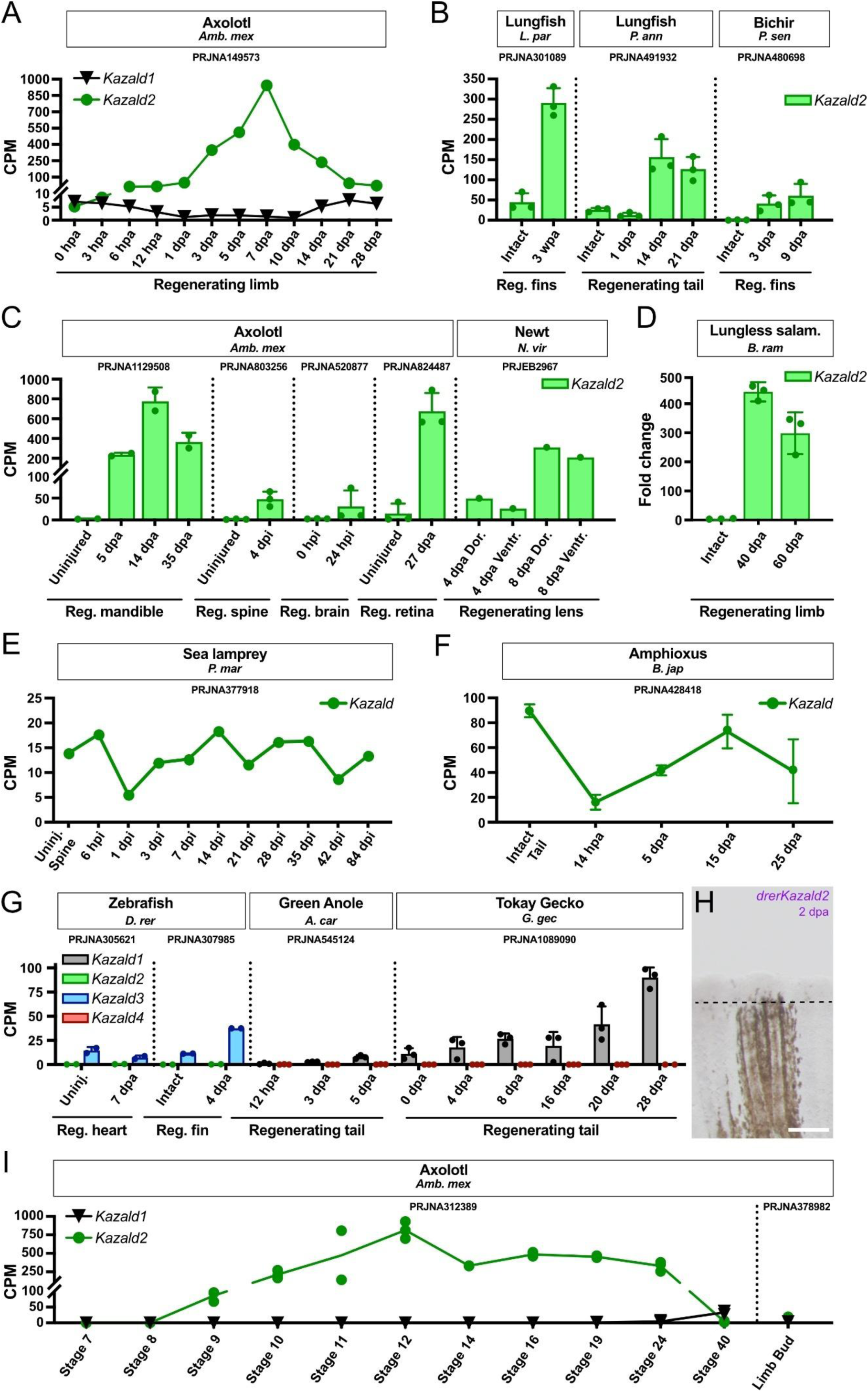
*Kazald2* is expressed during regeneration in species throughout the osteichthyan lineage. A) Quantification of *Kazald1* and *Kazald2* expression over the course of axolotl limb regeneration. B) Quantification of *Kazald2* expression in the regenerating fins and tail of lungfish and bichir. C) Quantification of *Kazald2* expression in the regenerating mandible, spine, brain, and retina of axolotl, and the regenerating lens of newt. D) RT-qPCR examining *Kazald2* expression in the regenerating limbs of lungless salamander. E) Quantification of *Kazald* expression over the course of lamprey spine regeneration. F) Quantification of *Kazald* expression over the course of amphioxus tail regeneration. G) Quantification of *kazald2* and *kazald3* expression in the regenerating heart and caudal fin of zebrafish, and *Kazald1* and *Kazald4* expression in the regenerating tails of green anole and tokay gecko. H) Whole mount *in situ* hybridization of *kazald2* in regenerating caudal fin of adult zebrafish at 2 dpa. Dashed line indicates the amputation planes. Scale bar: 250 μm. I) Quantification of *Kazald1* and *Kazald2* expression over the course of axolotl embryo development, and in the developing limb bud of axolotl larvae. PRJ IDs indicate the publicly available RNA-Seq datasets which generated the raw data for the listed tissues. Dots represent biological replicates in examined datasets, error bars represent standard deviation when calculable. CPM = Counts Per Million, hpa = hours post amputation, dpa = days post amputation, wpa = weeks post amputation, hpi = hours post injury, dpi = days post injury, Reg. = regenerating, Uninj. = uninjured, Dor. = dorsal, Ventr. = ventral.

The above-mentioned salamanders, lungfish, and bichir all possess a larval stage, after which most undergo metamorphosis into adulthood (70–72). We wondered if this extended post-embryonic development or maintenance of larval features may enable these species to utilize similar methods of regeneration, such as reactivating developmental pathways that are not restricted to embryonic stages. Thus, we measured *Kazald2* expression via RT-qPCR in regenerating limbs of the direct-developing lungless salamander *Bolitoglossa ramosi* (73) (**Fig. 6D**). This revealed a similarly large upregulation in the blastema even within a species that is neither paedomorphic nor undergoes metamorphosis.

As a connection of *Kazald2* to regeneration appears independent of the presence of calcified tissue, it could be that a Kazald gene has had this association since before the origin of bony vertebrates. However, since regeneration in cartilaginous fish is extremely limited, we could not determine if this connection existed in the jawed vertebrate ancestor. Thus, we looked further back in evolutionary time by investigating published RNA-Seq data of spine regeneration in the lamprey *Petromyzon marinus*, a jawless vertebrate. Its singular *Kazald* gene did not undergo large or consistent expression changes during regeneration, revealing that this process in lamprey likely occurs without use of a Kazald gene (**Fig. 6E**). Analysis of data from the even more distantly related amphioxus *Branchiostoma japonicum* found that its singular *Kazald* gene was strongly downregulated from high basal levels until near the end of regeneration, demonstrating that this gene is not positively correlated with this process (**Fig. 6F**). Finally, the one species of planaria we found that possesses a *Kazald* gene did not express it at detectable levels in data from either intact or regenerating tissue. Thus, it appears likely that there is an ancestral connection of *Kazald2* specifically with regeneration that evolved within the lineage leading to bony vertebrates following their split from cyclostomes, but it is unclear if this occurred before or after their split from cartilaginous fish.

However, there is one lineage of bony vertebrates with species that still possess *Kazald2* but do not express it during regeneration: teleost fish. Analysis of published RNA-Seq data of intact and regenerating fin and heart of zebrafish revealed almost no expression of *kazald2*, which we confirmed in regenerating fins at 2 dpa via WISH (**Fig. 6G and H**). Furthermore, regeneration occurs in teleost species that have lost *Kazald2*, such as medaka (74, 75). A possible explanation is that a different Kazald gene is expressed during regeneration instead, which is supported by the large expression of *kazald3* in the blastema and non-bone, inter-ray area of regenerating zebrafish fin (**Fig. 5D**). However, if this is the case, then *Kazald3* in teleost regeneration seems to be more tissue restricted than *Kazald2* in salamanders, as *kazald3* is not upregulated during zebrafish heart regeneration (**Fig. 6G**). Thus, another possibility is regeneration in teleosts evolved to not need a Kazald gene.

Such a development of Kazald gene-independent regeneration has seemingly occurred in lizards, as we found that the expression of neither of their Kazald genes peaked during the period of blastema development and growth. Analysis of published RNA-Seq data of tail regeneration in two lizard species found that *Kazald4* was mostly non-existent, and that while *Kazald1* continuously increased, its greatest expression in gecko occurred after the period of blastema formation and proliferation (**Fig. 6G**). Thus, its expression pattern better matches the growth of the replacement cartilage rod (76), which likely reflects the connection to skeletogenesis we have demonstrated. However, as the tail tissue in that study was always taken from a small region immediately distal to the amputation site (76), it could be that *Kazald1* is also expressed by undifferentiated cells in the blastema, but later work is needed to investigate this.

Finally, while we have shown that *Kazald2* possesses an ancestral connection to regeneration in at least the bony vertebrate lineage, the question remained if it could also be used in other processes. Part of the initial interest in this gene was due to its expression during axolotl limb regeneration, but not during limb development (9). Thus, it seemed to not be a developmental gene that was reactivated during regeneration, but instead a regeneration-specific gene. However, our analysis of published RNA-Seq data of axolotl embryonic development uncovered that although *Kazald2* is not expressed in the limb, it is highly expressed from gastrulation (stage 8) to the tailbud stage (stage 24) (**Fig. 6I**).

This window of expression aligns with its ortholog in the frog *Xenopus laevis*, which has been named *Mixer inducible gene 30 (Mig30)* (77). In contrast to axolotl though, the ortholog of *Kazald1* in *X. laevis* (labeled in that publication as xIGFBP-rP10), is also found in development. The lack of *Kazald1* for much of axolotl development also creates an inverse correlation between it and *Kazald2*, opening the possibility that their interplay could signal a transition between differentiated and undifferentiated cell states in this species. Finally, these genes serve as further examples of differences in gene names between animal models, hindering cross-species research if gene orthology is not easily accessible.

## Discussion

Through our work, we have uncovered the existence of the previously unidentified Kazald gene family. Furthermore, we have shown that this gene family is ancestral to all modern jawed vertebrates by tracing its origin to the two-round whole genome duplication event. Finally, we have investigated putative roles for each of the four genes of this family, and have determined when these roles most likely arose. Uncharacterized genes and genes with no published descriptions of their roles or functions, such as *Kazald4* and *Kazald3* respectively, are often overlooked in scientific research due to the large difficulties incurred by the lack of annotation and prior existing information (78). Meanwhile, the other two Kazald genes, *Kazald1* and *Kazald2*, have had their descriptions limited to just a few species. Thus, our work provides a valuable resource for both identifying what processes and/or tissues these genes are connected to in a much wider array of species, and for increasing the predictive power of how translatable their roles may be across species by determining how strongly a role has been conserved over evolutionary time.

### Kazald1 has an ancestral role in skeletogenesis and odontogenesis

The evolution of a mineralized skeleton and other skeletal elements, such as teeth and scales, is one of the most important developments within the jawed vertebrate lineage (79, 80), and thus fittingly serves as the basis for the name of the superclass we belong to: Osteichthyes. Exactly how this mineralized skeleton develops has been studied in a wide variety of distantly related species, including the mouse, chicken, salamander, and fish (52, 81, 82). Our findings contribute to this cross-species analysis by discovering that *Kazald1* has a connection to both skeletogenesis and odontogenesis across tetrapods. This strongly conserved association of *Kazald1* to this process implies that this gene and the molecular machinery surrounding it are likely to be under heavy selective pressure to remain stable. A probable conclusion is thus that the published findings about the functions of *Kazald1*, as well as the cell types expressing it, in mammals (21, 83) would also be applicable across tetrapods as a whole. However, future work to test this hypothesis is still required.

The expression of *Kazald1* in odontogenesis is likely even older than the tetrapod lineage, as our work identified a conserved expression of it not only in the teeth of the alligator and axolotl, but also in species of sharks. This greatly expands the connection between this gene and the teeth from its initial description in mice to now cover the entirety of jawed vertebrates. The most parsimonious explanation for this is that *Kazald1* has had a role in odontogenesis since before the split of the osteichthyan and chondrichthyan lineages, which would make it one of the most ancestral expression domains of any of the four Kazald genes. However, functional examination of how *Kazald1* is being utilized in the teeth of these different species is still required to determine if the exact role and function of this gene is equally as conserved.

Meanwhile, our examination of the connection of *Kazald1* to skeletogenesis shows that its expression in non-teleost actinopterygian fish is notably reduced in comparison to tetrapods and other sarcopterygians, and that it had been completely lost in the teleost fish lineage. The low expression in the bichir and paddlefish may be the result of their reduced, and overall cartilaginous, skeletons, which are thought to be derived traits (54, 57). Alternatively, the lack of expression, as well as the eventual loss in teleosts, may be due to the bony vertebrate ancestor not actually utilizing *Kazald1* in bone development. Indeed, various key differences in osteoblast gene expression between sarcopterygians and actinopterygians have indicated that certain osteoblast markers likely only developed following the split of these lineages (84). A lack of a firmly established role in ossification would also help explain its varied expression in some species of non-tetrapod/teleost fish, such as *Kazald1b* in paddlefish muscle and *Kazald1* in multiple different lungfish tissues. Distinguishing between these two possibilities will require work specifically focused on characterizing the expression profiles of osteoblasts in these actinopterygian and sarcopterygian fish in order to determine when *Kazald1* gained a role in skeletogenesis.

### Kazald2 has an ancestral role in gnathostome regeneration

The other Kazald with a presumed deeply ancestral role within the jawed vertebrate lineage is *Kazald2*, which we have shown to be connected to regeneration since at least the last common ancestor of extant osteichthyans. Since appendage regeneration is generally considered to be an ancestral trait that amniotes subsequently mostly lost (8), there is hope that findings in other species will one day be translatable to human patients. However, this idea is complicated not just by the vast differences that exist between mammals and the regeneratively-capable species currently studied, but also from findings that the mechanisms of regeneration can substantially differ even between more closely related species.

A key example is caudal fin regeneration in zebrafish vs. killifish, in which only a small fraction of differentially expressed genes is shared between the two species (85). These species diverged from each other approximately 230 million years ago (mya), much more recently than when humans diverged from zebrafish (∼430 mya), or even from salamanders (∼350 mya) (86). The larger amount of evolutionary time between humans and these lineages would suggest that even more differences have accrued between them, likely preventing many regeneration-associated genes from functioning the same in the two species. One way to help address this difficulty is by determining just how ancestral and strongly conserved the roles of individual genes are in a given process.

Presumably, if a gene has had a long-standing role across lineages, then the proteins and other molecules that it interacts with to perform that role are also likely to have been similarly conserved (87). The ability of a gene to interact with these other molecules may even outlive the gene itself, potentially by it being replaced by a non-orthologous gene or by the other molecules still retaining the interaction site (88, 89). We have shown that *Kazald2* upregulation has been most likely linked to regeneration since the bony vertebrate ancestor, and has been conserved in species from both the sarcopterygian and actinopterygian lineages. Thus, this indicates that there is a greater chance that its expression in non-regenerative species could induce some regenerative effects than if this role in regeneration had been a novel adaptation of the salamander lineage. The loss of *Kazald2* in tetrapods is also correlated with a vast reduction in regenerative potential, although it cannot be determined which of the two might have been the driving force for the other. Finally, the lack of *Kazald2* expression during regeneration in teleost fish like the zebrafish and medaka raises the interesting question of if these species are using entirely different pathways for this process than other bony vertebrates, or if they have replaced *Kazald2* with another gene, such as *Kazald3*.

### Kazald3 has evolved novel roles in teleost fish, but is commonly lost in other lineages

While the previous two Kazald genes both possess deeply ancestral expression profiles within the jawed vertebrate lineage, *Kazald3* does not. In fact, most tetrapod lineages have lost this gene, and in the few that still possess it, we were unable to identify a tissue or biological process in which it was expressed above very low levels. Not even the lungfish, which had the widest diversity of expressed Kazald genes across different tissues of any of the examined sarcopterygians, ever expressed *Kazald3* beyond extremely low levels. While a usage for *Kazald3* in sarcopterygians may exist that we were unable to identify, the lack of a role for it would explain the common loss of this gene in this lineage. Similarly, the frequent loss of *Kazald3* in many tetrapod lineages also supports the hypothesis that this gene has become decoupled from any biological tissue or process.

While *Kazald3* is commonly lost in sarcopterygians, we discovered that it is actually the most conserved Kazald gene in teleost fish of the actinopterygian lineage. Teleost fish constitute ∼96% of all fish species, and ∼50% of all vertebrates (90), and every one of its species that we analyzed possessed *Kazald3*. This remarkably strong maintenance across such a large and diverse clade implies that it has gained an important and overall conserved role within these species, which we have shown could be, at least partially, related to skeletogenesis.

Interestingly, this process is what *Kazald1* is connected to in other jawed vertebrates, and thus the association of *Kazald3* to this process in teleost fish would likely explain the highly unusual loss of *Kazald1* in this lineage. However, further work in other teleost fish species is important to verify the extent of this putative neofunctionalization, as certain species like the eel express multiple Kazald genes to a similar extent within their bones.

In addition to the overlap that we observed between *Kazald3* and the newly produced bone of regenerating zebrafish fin rays, we also found it to be expressed in a large surrounding region of non-skeletal tissue. This suggests that, in addition to taking on an expression profile similar to *Kazald1* in other lineages, it may potentially have also replaced the use of *Kazald2* within regeneration. This is further supported by our findings that *Kazald2* is not upregulated during zebrafish regeneration, and that this gene has been completely lost in medaka without impairing its ability to regenerate the fin (74, 75). However, this requires further testing, as the lack of clear upregulation in the zebrafish regenerating heart indicates that its usage may not be as widespread as that of *Kazald2* in salamanders.

### Kazald4 is highly conserved in gnathostomes without a clear use, but has evolved a novel role in avian brains

*Kazald4* represents an interesting contrast to *Kazald3*, as it appears to be the second most conserved Kazald gene in jawed vertebrates, when measured by the number of orders still possessing it, but it does not appear to have a correspondingly strongly conserved role. From our examined sarcopterygians, the lungfish did display moderate to strong expression of *Kazald4* in various tissues, such as the lung, jaw, skin, and heart. However, this wide range of expression was not reflected in tetrapods, with most of the species we analyzed not expressing it above low levels in any of the examined tissues or biological processes. This was also the case for actinopterygians, as non-teleost fish, like the bichir and paddlefish, had very low to nonexistent expression in all of their different tissues. Meanwhile, one teleost fish, the eel, did weakly express *Kazald4* in its bone, but other Kazald genes were also expressed in this tissue at similar levels. Unfortunately, additional analysis in teleost fish was impaired by the most common model species, zebrafish and medaka, not possessing *Kazald4*. Therefore, further examination of other teleost fish that still maintain this gene is needed to discover if a clear role for *Kazald4* may exist in at least a part of this diverse lineage.

Despite this lack of a clear and conserved role across the majority of bony vertebrate lineages, we found that birds appear to have developed a strong use for it. Distantly related avian species all expressed high levels of *Kazald4* throughout the brain, which was not observed in any other tetrapod. While our work cannot directly answer what the exact function of this gene in the avian brain is, the high similarity of the four Kazald genes at a genetic and proteomic level makes it plausible that they may perform similar functions at a cellular level. Mainly *Kazald1* has been functionally characterized and found to promote cell proliferation (83). Therefore, it is feasible that *Kazald4* is also involved in cell proliferation in the avian brain. Such an idea is particularly intriguing as, unlike mammals in which adult neurogenesis is limited, birds continue to generate new neurons throughout their lives, with some regions even being described as “hot spots” of neuronal proliferation (91, 92). Thus, future studies on if the centers of *Kazald4* expression correspond to these areas, and especially on what impact its inhibition has on the brain, might provide exciting insights into adult neurogenesis in birds.

As birds are the only lineage of bony vertebrates we could find in which *Kazald4* was highly expressed, its strong conservation across jawed vertebrates is perplexing. Without an apparent function in many of these lineages that would drive purifying selection, it appears strange that it has not been lost as often as many of the other Kazald genes. While a potential explanation is that there actually is a conserved function for *Kazald4* that we were unable to uncover, another fascinating possibility is that the characteristics of the genomic region containing *Kazald4* may be contributing to the preservation of this gene. Several genomic features have been associated to elevated rates of gene loss, including high GC content, high gene density, high densities of repetitive elements, and high nucleotide substitution rates (93, 94). Thus, if over the course of jawed vertebrate evolution, the genomic regions containing *Kazald4* have consistently lacked these features in comparison to the regions containing the other Kazald genes, then it may not have been as susceptible to loss. To address what constitutes high gene density or substitution rates in such distantly related species is complicated, since the genomes can differ in size by orders of magnitude and can be changing at very different rates. However, future work examining these genomic regions may be able to use this information to predict if *Kazald4* is likely to possess a functional role we were unable to uncover, or if its persistence in vertebrate genomes could be due to factors beyond the gene itself.

### Possibility for specific Kazald gene replacement with paralogs

Potentially the most interesting question remaining is if specific Kazald genes are able to act in place of their paralogs. Our work has found that all four Kazald genes possess the same protein domains in the same order, and that the three-dimensional structure of the regions containing these domains are quite similar to each other. Such a high similarity could indicate that the different Kazald paralogs are able to functionally substitute for each other, which is supported by our finding that *Kazald3* appears to have taken over the ancestral role of *Kazald1* in teleost fish. Therefore, future work will hopefully be able to examine the possibility that at least certain Kazald genes could act in place of each other.

## Materials and Methods

### Gene nomenclature usage

Published nomenclature guidelines were followed when referencing gene names and symbols within specific species, such as axolotl (e.g., *Kazald1*) (95), mouse (e.g., *Kazald1*) (96), zebrafish (e.g., *kazald3*) (97), and chicken (e.g., *KAZALD1*) (98). When referencing other species or when discussing genes outside of a species-specific context, the guidelines of the axolotl and mouse were followed.

### Kazald gene identification in axolotl and other species

The current axolotl (*Ambystoma mexicanum*) genome and transcriptome (RefSeq assembly GCF_040938575.1; WGS project JBEBLI01) were searched through using TBLASTN (version 2.14.1 (99)) run via command line with default parameters using the amino acid sequences of mouse (*Mus musculus*) *Kazald1* and zebrafish (*Danio rerio*) *kazald2* and *kazald3*. The genomes and/or transcriptomes of all other used animal species were downloaded and initially searched through for potential Kazald genes using reciprocal TBLASTN with the amino acid sequences of mouse *Kazald1*, zebrafish *kazald2* and *kazald3*, and the four putative axolotl Kazald genes (**Table 1**). Animal genomes were chosen to extensively cover the jawed vertebrate lineage, and to provide representatives from major lineages of jawless vertebrates, invertebrate deuterostomes, protostomes, and non-bilaterian animals. Annotated genomes available from NCBI were preferred for selection, but was not a requirement.

### Identification of conserved features of putative Kazald genes

The four-exon structure that we considered to be characteristic for vertebrate Kazald genes (**Supp. Fig. 3**) and used for subsequent procedures, was identified by checking the reported exon-intron structure of putative Kazald genes in the gff annotation files of examined species. Three protein domains arranged in a specific order were considered to be characteristic of all Kazald genes, which were: (1) Insulin-like growth factor-binding protein domain (Igfbp domain); (2) Kazal-type serine protease inhibitor and follistatin-like domain (Kazal domain); (3) Immunoglobulin-like domain (Ig-like domain). This protein domain order was used for subsequent procedures, and was identified via the Expasy webserver ScanProsite tool (100) using the putative Kazald gene amino acid sequences listed in available transcriptomes.

### Identification of unannotated Kazald genes in genomes of species

The genomes of all examined species, when available, were searched through for potential Kazald genes that had not been annotated. This was done in two steps:

1. The genome was searched via TBLASTN using the putative Kazald gene amino acid sequences of the axolotl, gray bichir (*Polypterus senegalus*), West African lungfish (*Protopterus annectens*), and several closely related species to the investigated species, along with the Kazald genes provided in the transcriptome of the investigated species when available. The genome was also searched with the amino acid sequence of the gene *insulin-like growth factor binding protein 7* (*Igfbp7*) of the axolotl and the investigated species, when available, as a control. If there were genomic locations that were more similar to a Kazald gene than they were to *Igfbp7*, and which did not contain any annotated Kazald genes, then these locations were analyzed in Step 2.
2. The identified locations were analyzed via Exonerate (version 2.2.0 (101)) and Miniprot (version 0.12-r237 (102)) using the amino acid sequence of its best match Kazald gene within each of the species used for the TBLASTN search of Step 1. Both programs were run with relaxed parameters, which were modified on a case-by-case basis, to determine if there was a gene sequence in the examined location that could be a Kazald gene. In vertebrates, the sequence was checked for possession of the characteristic four-exon structure and protein domain order, while in invertebrates only the protein domain order was checked using the Expasy webserver ScanProsite tool. Sequences that fulfilled these criteria were then searched for internal stop codons or frameshift mutations in the Exonerate/Miniprot output files. If none were found, then it was identified as an unannotated Kazald gene that was used in future analysis. Otherwise, the sequence was categorized as a potential pseudogene.

### Editing of annotated Kazald genes

If an annotated Kazald gene within a species was highly dissimilar from the Kazald genes of closely related species, then it was examined to see if the annotation could be incorrect. In vertebrates, this was also done if the annotated gene did not possess the characteristic four-exon structure. Examination was done through two steps:

1. The genomic location containing the dissimilar Kazald gene was analyzed via Exonerate and Miniprot using default parameters with the Kazald genes of closely related species. In vertebrates, the used Kazald genes had to have the characteristic four-exon structure and protein domain order. If a similar sequence to these Kazald genes was found, then the analysis proceeded to Step 2.
2. The genomic locations within the closely related species that contained the Kazald genes used for Step 1 were reciprocally analyzed via Exonerate and Miniprot using default parameters with the dissimilar Kazald gene from Step 1.

If Step 1 uncovered a sequence that was highly similar to the Kazald genes of the closely related species, then that sequence was identified as a putative isoform of the dissimilar Kazald gene, and replaced the dissimilar Kazald gene for use in subsequent analyses. Additionally, if Step 2 failed to discover a sequence in any of the closely related species that was highly similar to the dissimilar Kazald gene, then the dissimilar Kazald gene was determined to likely be an incorrect annotation.

### Phylogenetic analysis with maximum likelihood

Maximum likelihood phylogenetic trees were created from the whole amino acid sequences of putative Kazald genes using RAxML-NG (version 1.2.0 (103)). The peptide sequences were first aligned using MAFFT (version 7.520 (104)), with the L-INS-i alignment setting and default parameters. The JTT+I+G4 or JTT+R4 substitution models were used depending on the gene, which were chosen via ModelTest-NG (version 0.1.7 (105, 106)). RAxML-NG was run based on commands described in the tutorial page of the program wiki (github.com/amkozlov/raxml-ng/wiki/Tutorial). A convergence cutoff of 1% was set to determine the number of bootstraps to run.

### Phylogenetic analysis with Bayesian inference

Bayesian inference phylogenetic trees were created from the whole amino acid sequences of putative Kazald genes using BAli-Phy (version 3.6.0 (107)). BAli-Phy was run using the following commands based on the type of analysis.

Analysis of Kazald gene family in all species (6 chains were run): `bali-phy Kazald_Peptides.fa -S jtt+Rates.free[n=4] --set infer-ambiguous-observed=true -n Kazald`.

Analysis of one specific gene family in limited species (4 chains were run): `bali-phy Gene_Peptides.fa -S jtt+Rates.free[n=4] --set infer-ambiguous-observed=true -n GeneName`.

Analysis of the syntenic blocks of limited species through partitioned analysis (4 chains were run): `bali-phy Adra1_Peptides.fa FgfD_Peptides.fa Gfra_Peptides.fa Kazald_Peptides.fa Lrrtm_Peptides.fa -S jtt+Rates.free[n=4] -n SyntenicBlocks`.

The JTT+R4 substitution model was used for all runs, as it was chosen as either the best or second-best model for all genes via ModelTest-NG. BAli-Phy was run until the chains converged, which was determined using the included bp-analyze script, and the program Tracer (version 1.7.2 (108)).

### RNA-Seq read mapping and expression analysis

Downloaded FASTQ files from analyzed BioProjects were trimmed of adapter sequences and low-quality bases using the programs cutadapt (109) and fastq_quality_filter from the FASTX-Toolkit (https://github.com/agordon/fastx_toolkit), respectively.

If a genome was available, then the reads were then mapped against it via the program HISAT2 (version 2.2.1 (110)). HISAT2 was run through the command line with standard default parameters and a known-splicesite-infile created from the gff annotation file via the hisat2_extract_splice_sites.py script. Transcript quantification was conducted using StringTie (version 2.2.1 (111, 112)) through the command line with standard parameters and the option of assembling novel transcripts. Finally, normalized counts per million (CPM) values for each sample were calculated using the Bioconductor package edgeR (version 3.40.2 (113)), for R (version 4.2.2 (R Core Team, 2021) (R-project.org)).

If a genome was not available, then the reads were mapped against the transcriptome via the program Salmon (version 1.10.1 (114)). Salmon was run through the command line with default parameters. Finally, normalized CPM values for each sample were calculated using the Bioconductor package edgeR for R.

### Kazald protein 3D structure prediction

Kazald proteins had their 3D structure predicted with AlphaFold2 through the use of the online tool ColabFold: AlphaFold2 using MMseqs2 (version 1.5.5) hosted on a Google Colaboratory Notebook (115, 116). Amino acid sequences were used as the query and the tool was run with default parameters.

### Kazald protein 3D structure alignment and similarity quantification

Kazald proteins were aligned to each other and had their similarity quantified using the Universal Structural alignment (US-align) program through the online webserver hosted by the Zhang lab (117). The structural information of proteins, provided within Protein Data Bank (.pdb) files, were uploaded and the tool was run with default parameters.

### Kazald protein 3D structure visualization

The 3D structure of both individual and aligned Kazald proteins were visualized using the Mol* 3D Viewer hosted by the RCSB Protein Data Bank (118, 119). Visualization was made using the structural information of proteins provided within Protein Data Bank (.pdb) files.

### Animal husbandry

Axolotl husbandry and experimental procedures were performed according to the Animal Ethics Committee of the State of Saxony, Germany. Animals used were selected by their size (snout-to-tail and snout-to-vent lengths). Husbandry was performed in the Center for Regenerative Therapies Dresden axolotl facility using methodology adapted from (120) and according to the European Directive 2010/63/EU, Annex III, Table 9.1. Axolotls were kept in conditions described in (16).

Zebrafish husbandry and experimental procedures were performed according to the animal handling and research regulations of the Landesdirektion Sachsen, Germany (permit numbers: DD24.1-5131/450/4 and 25-5131/564/2 and respective amendments). Husbandry was performed as described in (121). Adult zebrafish of both sexes were used. The transgenic fish line *runx2*:GFP=Tg(*Hsa.RUNX2-Mmu.Fos*:EGFP)^zf259^ has been described (68).

Fertilized eggs of the small-spotted catshark (*Scyliorhinus canicula*) were incubated to harvest developing embryos, complying with the guideline defined by the Institutional Animal Care and Use Committee at National Institute of Genetics (Approval ID: R5-14 and R6-13). Embryos were staged according to the developmental stages established in (122).

Wild adult (7-10 cm, snout-to-tail length) *Bolitoglossa ramosi* lungless salamanders were collected from their type locality in the Andes region of Antioquia, Colombia, under the Ministerio del Medio Ambiente contract on access to genetic resources number 118−2015. All experimental procedures were approved by the Institutional Animal Care and Use Committee of the University of Antioquia. These salamanders were collected and kept as described in (123).

### Axolotl surgery and tissue collection

Axolotls were anesthetized with 0.01% benzocaine solution (Sigma-Aldrich, #E1501-100G) by immersion. Amputations were performed with a scalpel through the forelimb at the mid-radius/ulna level. Following amputation, animals were kept on benzocaine for 15 minutes and then transferred back to water and allowed to regenerate at 20°C.

Tissue collection was performed by euthanizing animals in a lethal dosage of 0.1% benzocaine solution by immersion for at least 20 minutes. For HCR, whole stage 46 embryos (49, 51) were collected and fixed in 4% formaldehyde in 1× phosphate buffered saline (PBS) for 40–60 minutes and stored in 100% ethanol at −20 °C. For paraffin sectioning and embedding of 15 days post-amputation (dpa) limbs, tissue from three 6 cm (snout-to-tail) animals was fixed in 1× MEMFa (0.1 M MOPS pH 7.4, 2 mM EGTA, 1 mM MgSO4·7H2O, and 3.7% formaldehyde) for a minimum of 3 days, and then decalcified in RNase-free 0.5 M EDTA for 1 week with daily changes of solution. Tissue collection and fixation of adult mandible tissue was performed as in (16).

### Zebrafish fin clips and tissue fixation

Zebrafish were anesthetized with 0.02% Tricaine (Sigma-Aldrich, #A5040) by immersion. Fin clips were performed at 50% of fin length with a scalpel. Animals were transferred to fish water and allowed to regenerate at 28°C.

Fin regenerates and embryos were fixed in 4% paraformaldehyde (PFA) in PBS overnight at 4°C at the indicated times post fin clip and stage, respectively. After fixation, fins and embryos were washed in PBS and dehydrated in methanol for storage at −20°C.

### Scyliorhinus canicula embryo collection

Embryos were extracted from their egg cases by removing the surface layers with a knife and forceps. Embryos were fixed in 4% PFA in PBS overnight at 4°C. After fixation, embryos were dehydrated in PBS/methanol, and stored in 100% methanol at −20°C.

### Bolitoglossa ramosi surgery and tissue collection

Animals were anesthetized with 1% tricaine (Sigma-Aldrich, #E10521) by immersion. Amputations were performed as described in (123). Briefly, animals were placed in a Petri dish containing 20 mL tricaine for 4 minutes. Amputations were performed with microscissors through the forelimb at the mid-humerus level. Protruding bone and muscle were trimmed to obtain a flat wound surface. Following amputation, the wound was rinsed with 1 mL 0.5% sulfamerazine (Sigma-Aldrich, #S0800) to avoid infection. Animals were rinsed with abundant water to remove traces of tricaine, and were transferred to plastic containers and allowed to regenerate at 20°C.

Tissue collection was performed by euthanizing animals in a lethal dosage of 2% tricaine solution by immersion. Tissue was then collected and stored in TRIzol® reagent until total RNA was extracted following the reagent manufacturer’s protocol (Life Technologies).

### Paraffin sectioning and Movat’s pentachrome staining

Sample embedding, sectioning and staining of axolotl limbs and adult mandibles was performed by the CMCB Histology Facility, Dresden. Briefly, samples were dehydrated in a series of EtOH in RNase-free water until 100% EtOH, and then embedded in paraffin. Longitudinal sections of 4-5 µm were generated using a microtome. Movat’s Pentachrome (Morphisto Art-Nr:12057) staining in axolotl limbs was performed according to the manufacturer’s instructions. Imaging was performed using an Olympus OVK automated slide scanner system (UPLFLN 4x/0.13 or UPLSAPO 10x/0.40).

### Hybridization Chain Reaction (HCR) staining

Whole mount HCR was performed according to (124) with some modifications. Briefly, limbs were rehydrated through a series of MetOH in RNase-free water and washed three times in PBT (0.1% Tween 20 in PBS). Tissue was then delipidated in Delipidation Solution (200mM Boric acid, 4% SDS, pH 8.5 in RNAse-free water) for 2 hours at 37°C. After three washes in PBT, limbs were permeabilized with Permeabilization Solution (0.3M Glycine, 2% Triton X-100, 20% DMSO in PBS) for 1 hour at room temperature (RT). Limbs were washed again in PBT, incubated in pre-warmed Hybridization Buffer (Molecular Instruments, BPH01726) for 5 minutes and then pre-hybridized in new Hybridization Buffer for 30 minutes at 37°C. After this, tissue was incubated overnight with Hybridization Buffer containing 2 pmol per 500 µl of probe solution. The following day, limbs were washed four times with agitation for 15 minutes with Wash Buffer (Molecular Instruments, BPW01726) at 37°C and two times for 5 minutes in 5× SSCT (3M NaCl, 300 mM sodium citrate, 0.1% Tween 20, in water) at room temperature. Pre-amplification was performed for 5 minutes at RT in Amplification Buffer (Molecular Instruments, BAM01826), followed by amplification for 16-24 hours at RT in Amplification buffer with 30 pmol of each hairpin. Finally, tissue was extensively washed in 5× SSCT, incubated overnight with Hoechst 33258 (Abcam Ab228550) 1:1000 in PBS, and cleared in RIMS (LifeCanvas technologies, EI-500_1.52) for a minimum of one overnight. Regenerating zebrafish fins and stage 26 axolotl jaws dissected from the embryo were mounted in a glass bottom dish, and then imaged using a Zeiss LSM 980 inverted confocal laser scanning microscope (Plan-apochromat 10x/0.45).

HCR in slides was done according to (125). Briefly, slides were dewaxed in Roti-Histol (Carl Roth 6640) and rehydrated through a series of EtOH in RNase-free water. After washes in RNAse-free PBS, slides were treated with proteinase K (10 μg/ml in PBS) at 37°C for 10 minutes. Slides were then washed in RNAse-free water, moved into a humidified chamber containing a solution of 1:1 formamide and 2× SSCT, and pre-hybridized with pre-warmed Hybridization Buffer for 30 minutes at 37°C. Next, slides were drained from the pre-hybridization solution, covered in Hybridization Buffer containing the probes, protected from drying out with a glass coverslip and incubated overnight at 37°C in the humid chamber. In the following day, the coverslips were removed and the slides were sequentially washed for 15 minutes at 37°C with 100% HCR Wash Buffer, 75% Wash Buffer/ 5× SSCT, 50% Wash Buffer/ 5× SSCT, 25% Wash Buffer/ 5× SSCT, and 5× SSCT. The slides were then moved to a humidified chamber containing water and pre-amplified with Amplification Buffer for 30 minutes at RT. Next, slides were covered with a solution of Amplification Buffer containing snap cooled hairpins, covered with parafilm, and incubated for 24 hours in a dark humidified chamber at RT. Finally, the slides were extensively washed with 5× SSCT, incubated for 10 minutes with Hoechst 1:1000 in 5× SSCT, and mounted in VectaShield (Vector Laboratories, H-1000-10).

Probe sets for axolotl *Kazald1* (*amexKazald1*), zebrafish *kazald3* (*drerKazald3*), and *Gfp* were designed using the HCR probe generator created by the Monaghan Lab (https://github.com/Monaghan-Lab/probegenerator) (126) and purchased as oligo pools (O’Pools™ Oligo Pools) from IDT (Integrated DNA Technologies).

Each HCR was performed in a minimum of 3 different axolotl stage 46 embryos and zebrafish 4 dpa fin regenerates, or 3 different slides corresponding to 3 different animals in axolotl adult mandible tissue.

### Cloning of RNA probes for in situ hybridization

Probes for catshark *Kazald1* (*scanKazald1*), and zebrafish *kazald2* and *kazald3* (*drerKazald2* and *drerKazald3*) were amplified from genomic DNA using primers flanking the first exon of the corresponding gene. Each fragment was then cloned into pGEM-T easy vector system (Promega), according to the manufacturer’s instructions. Constructs were sequenced using the Mix2Seq Kit (Eurofins Genomics, Ebersberg, Germany) to select for inserts with the correct sequence. Prior to transcription, 10 μg of plasmid were linearized to obtain antisense and sense probes. For synthesizing RNA probes, *in vitro* transcription was carried out using a T7 polymerase (#RPOLT7-RO, Roche, Mannheim, Germany) or a SP6 polymerase (#RPOLSP6-RO, Roche), following manufacturer’s instructions.

### Whole mount in situ hybridization

Whole mount *in situ* hybridization (WISH) was performed using *in vitro* transcribed digoxigenin-labelled antisense RNA probes.

The protocol was adapted from (127) and performed as in (128). Before RNA *in situ* hybridization, samples were dehydrated to 100% MetOH with serial washes of MetOH in RNAse-free water and stored at −20°C. At the start of the protocol, samples were bleached in MetOH + 6% H2O2 at RT and fully rehydrated with decreasing concentrations of MetOH in TBST (1× TBS, 0.1% Tween 20) until 100% TBST. Tissues were then washed three times with TBST and treated with 10 μg/mL proteinase K (Pk) in TBST at 37°C. The timing of Pk treatment were as follows: whole mount stage 29 catshark embryos were incubated for 10 minutes, zebrafish embryos at 3 and 5 dpf were incubated for 15 minutes, and regenerating zebrafish fins for 20 minutes. After incubation, samples were washed with TBST and rinsed with 0.1M trietanolamine (Sigma-Aldrich, #90278) in RNAse-free water (TEA) pH 7.5. Tissue was next incubated with freshly prepared 0.1M TEA + 1% acetic anhydride (Sigma-Aldrich, #320102) for 10 minutes and then washed again with TBST. Next, samples were re-fixed with 4% PFA + 0.2% glutaraldehyde (Sigma-Aldrich, #G6257) for 20 minutes and washed with TBST. TBST was removed, and samples were incubated with previously warmed hybridization solution (50% formamide, 5× SSC [3 M NaCl, 300 mM sodium citrate] (pH 5.5), 0.1% Tween 20, 50 μg/ml yeast tRNA, 100 μg/ml heparin, 1× Denhart’s, 0.1% CHAPS, 5 mM EDTA) at 65°C for 4 hours. Tissue was incubated with Hybridization Solution containing the RNA probe overnight at 65°C and then washed at 65°C the following day twice with prewarmed 5× SSC (50% formamide, 5× SSC, 0.1% Tween 20), 2× SSC (50% formamide, 2× SSC, 0.1% Tween 20), and 0.2× SSC (0.2× SSC, 0.1% Tween 20) for 30 minutes each wash. Samples were washed with TNE buffer (10 mM Tris-HCl (pH 7.5), 500 mM NaCl, 1 mM EDTA), treated with RNase (20 μg/ml in TNE buffer) for 15 minutes, and washed again with TNE buffer. Next, tissue was equilibrated with MABT (100 mM Maleic acid, 150 mM NaCl, 0.1 % Tween 20), blocked with MABT/Block (MABT containing 1% blocking reagent (Roche, #11096176001)) for 1 hour at RT, and incubated with a 1:5000 dilution of alkaline phosphatase-conjugated anti-digoxigenin antibody (Roche, #11093274910) in MABT/Block overnight at 4°C. After extensive washes with MABT at RT, samples were equilibrated in NTMT (100 mM Tris-HCL (pH 9.5), 50 mM MgCl2, 100 mM NaCl, 0.1% Tween 20), and developed at RT in BM Purple (Roche, #11442074001). Reactions were stopped with PBS and fixed with 4% PFA overnight.

### RT-qPCR in B. ramosi

RNA was extracted from unamputated and regenerating (40 and 60 dpa) forelimb tissue samples from *B. ramosi* lungless salamanders using TRIzol® reagent (Life Technologies). RNA was then reverse-transcribed to single-stranded cDNA with reverse transcriptase (Thermo) in the presence of random hexamer primers, oligoDT primers, and dNTPs for 60 min at 42°C. Expression levels of specific mRNAs were determined by qPCR using gene-specific primer pairs, with three technical replicates. Each reaction was performed at a total volume of 10 μL containing 50 ng first-strand cDNA, 5 μL Syber greenMix (Biorad), and 0.1 μM of each primer pair, and cycled on a Biorad Real-Time PCR system. Real-time data were analyzed using Biorad software version 2.1. Relative mRNA expression was calculated using the 2 –ΔΔCT method with GAPDH as a cross-sample reference.

## Acknowledgments

We thank past and current members of the Sandoval-Guzmán lab for their support during the development of this work. We are also grateful to Anja Wagner, Beate Gruhl and Judith Konantz for their dedication to axolotl care. This work was supported by the Light Microscopy Facility and the Histology Facility, both Core Facilities of the CMCB Technology Platform at Technische Universität Dresden. SDK was supported by the PhD program of the DIGS-ILS and funding from the TU Dresden Graduate Academy. We thank Tomoyuki Satonaka at Shima Marineland, Kazuyuki Yamada at Tokai University Marine Science Museum, Daiki Katooka at Enoshima Aquarium, and Hatsune Makino-Itou at National Institute of Genetics, for catshark embryo sampling. The work at the TU Dresden is co-financed with tax revenues based on the budget agreed by the Saxon Landtag.

## Supplementary Figures and Tables

**Supplementary Figure 1.**
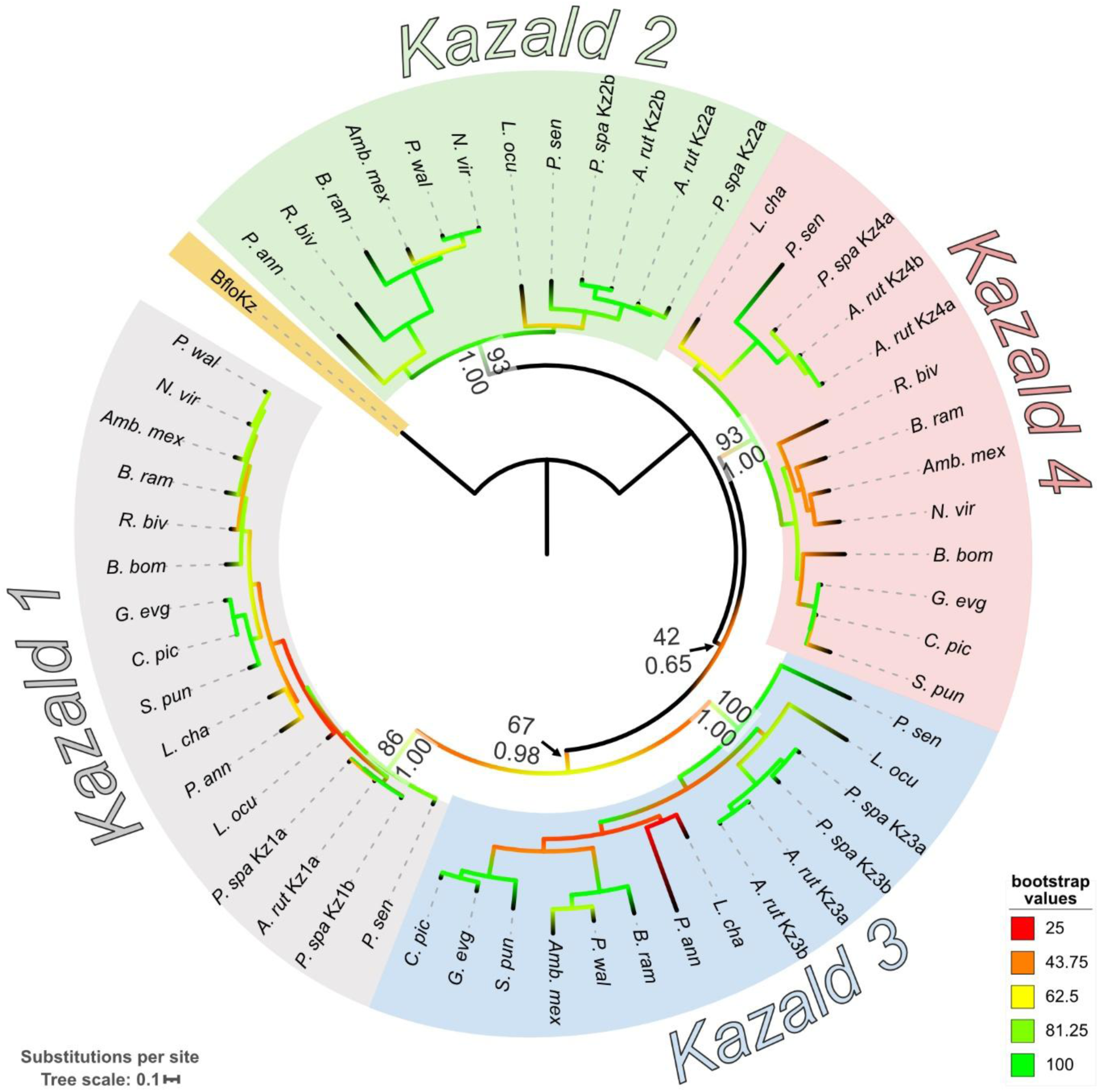
Phylogenetic tree of Kazald genes from a subset of species strengthens the support of the four distinct clades in vertebrates. Displayed consensus tree was generated via RAxML using amino acid sequences. Branch colors indicate the bootstrap values at each node. Support values are shown for the nodes basal to the four Kazald clades. Layout of the support values are: ML bootstrap support via RAxML (top), BI posterior probabilities via BAli-Phy (bottom). Abbreviated species names are listed in **Supplementary Table 2**. Kazald a and b versions created by whole genome duplications that occurred after the 2R-WGD event in the Acipenseriformes order of fish are marked next to the abbreviated species name. Scale bar corresponds to mean number of amino acid substitutions per site.

**Supplementary Figure 2.**
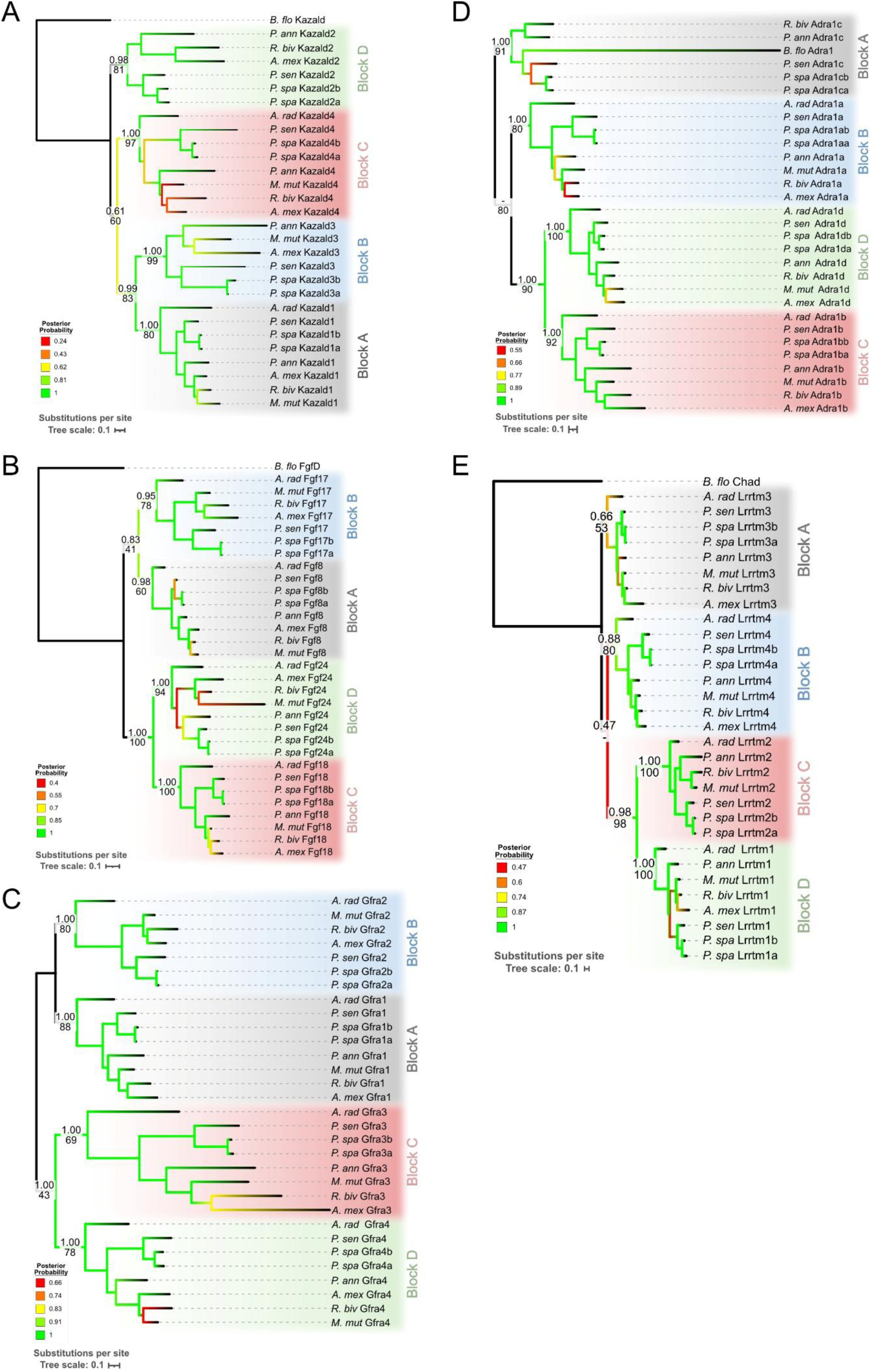
Bayesian inference trees of each of the multigene families chosen for the partitioned analysis of the Syntenic Blocks demonstrate no discordant topologies. Displayed trees were generated via BAli-Phy using amino acid sequences. Branch colors indicate the BI posterior probability at each node. Support values are shown for the nodes basal to the four Syntenic Blocks. Layout of the support values are: BI posterior probabilities via BAli-Phy (top), ML bootstrap support via RAxML (bottom). Scale bars correspond to mean number of amino acid substitutions per site.

**Supplementary Figure 3.**
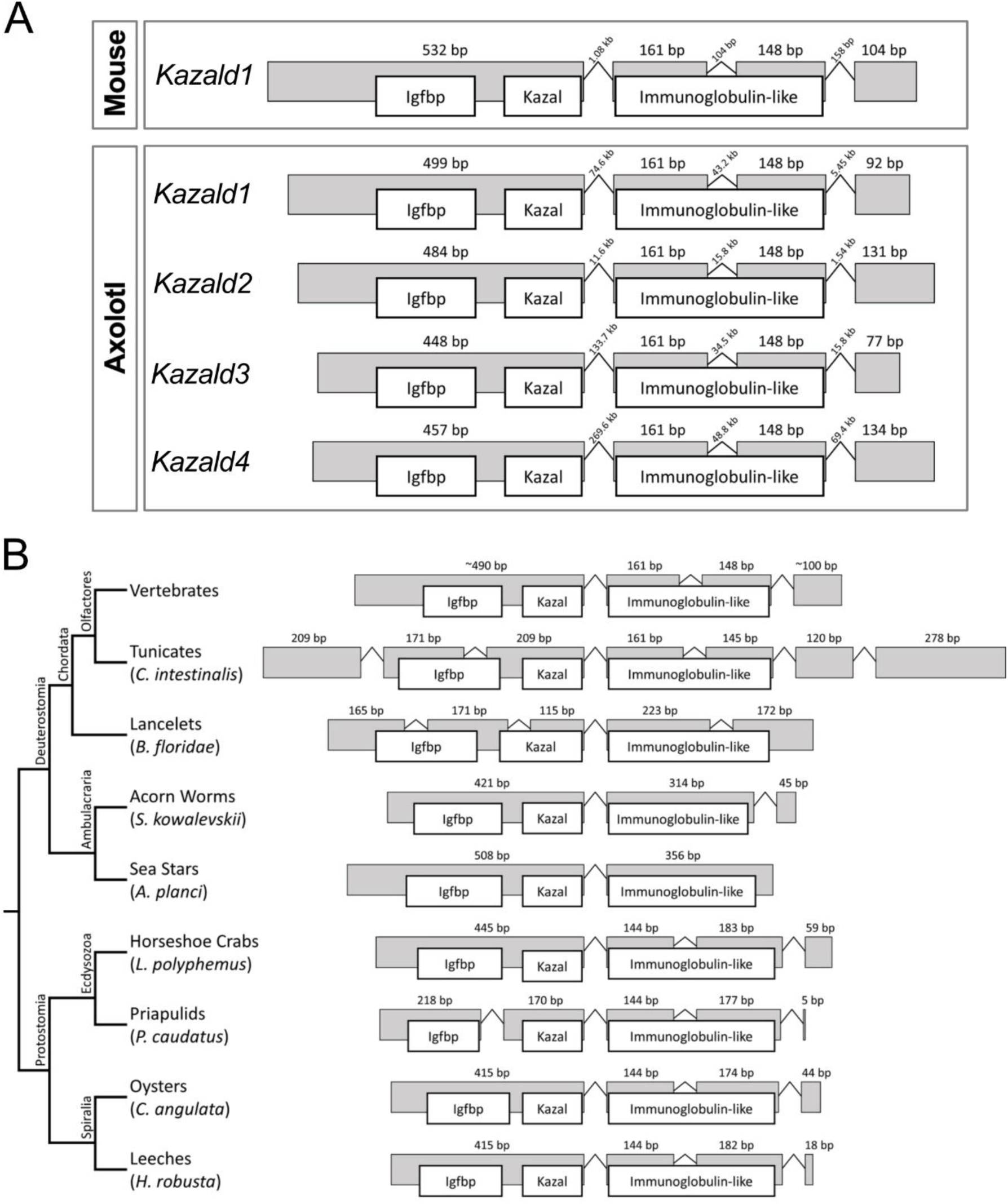
The four jawed vertebrate Kazald genes have a consistent genomic structure, which is also present in protostomes but not invertebrate deuterostomes. Exons are colored in gray and have their relative sizes scaled, introns are represented by bent lines and do not have their sizes scaled. Identity and location of the protein domains are represented by the overlapping white boxes, and are scaled to the size of the exons. A) Exon-intron structure and protein domain location comparison between mouse *Kazald1* and the four axolotl Kazald genes. B) Overview of the Kazald gene exon-intron structures and protein domain locations across bilaterians. Abbreviations: bp = base pairs, kb = kilobase.

**Supplementary Figure 4.**
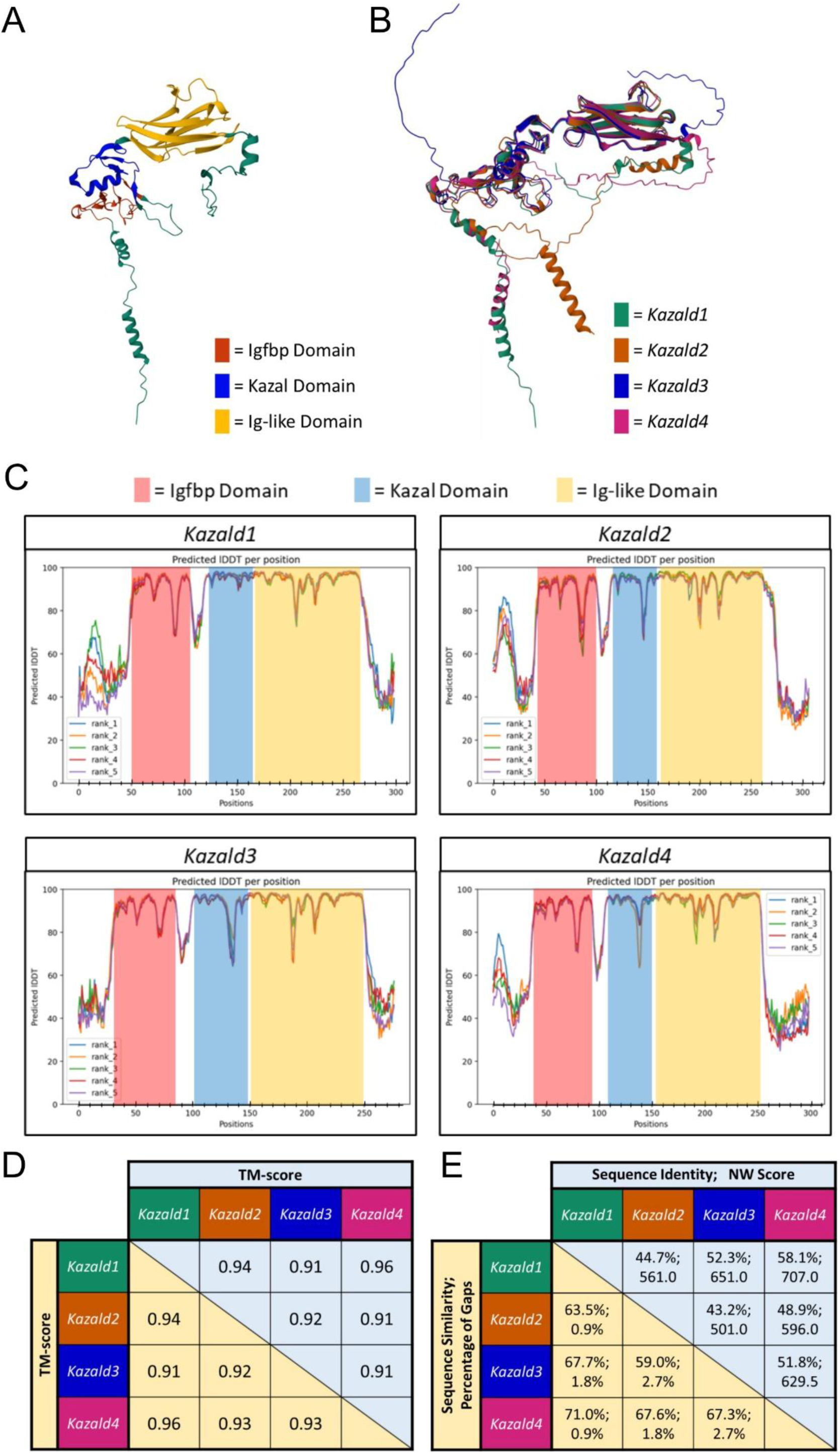
The regions of each Kazald gene that contain the three protein domains have similar 3D structures. A) Schematic of the protein structure of axolotl *Kazald1* with the three protein domains highlighted. B) Schematic of how the four axolotl Kazald proteins best overlap with each other. C) Graphs depicting the level of confidence that AlphaFold2 has in the predicted protein structure, expressed as the predicted local distance difference test (Predicted lDDT) per position. Five structures for each Kazald gene were predicted and then ranked through AlphaFold2, with the “rank_1” prediction of each one used for all subsequent analysis. D) Template modeling score (TM-score) of the superimposition of the Protein Domain Containing Regions (PDCRs) of the Kazald gene of the row against the Kazald gene of the column via US-align. As the TM-score is normalized by sequence length, the mirrored TM-scores can be different from each other. A TM-score of 1 equates to the two structures being 100% identical, a TM-score above 0.9 means the structures are nearly identical to one another (129), E) Amino acid sequence identity, sequence similarity, percentage of gaps, and NW Score of the PDCRs of two Kazald genes compared against each other via the Needleman–Wunsch algorithm.

**Supplementary Figure 5.**
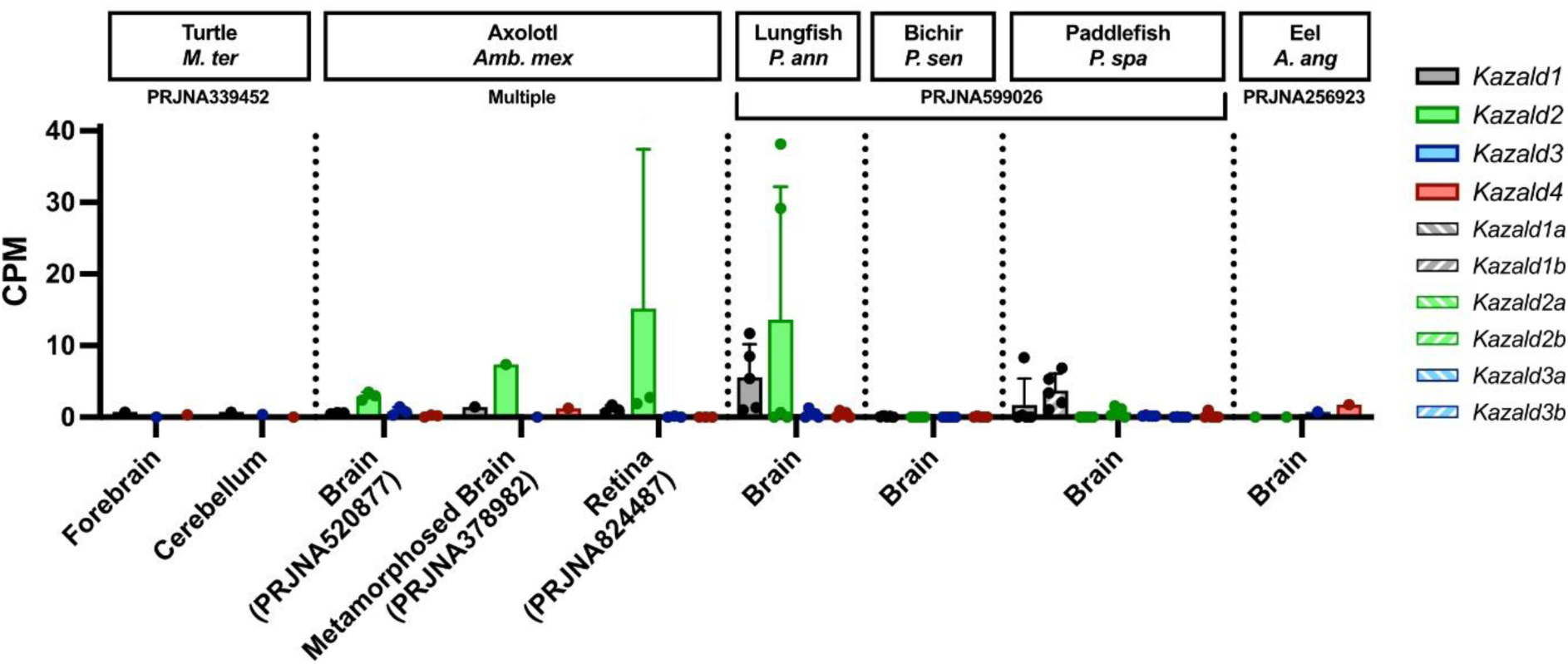
Non-avian species lack a strong and consistent expression of any Kazald gene in their brains. Quantification of the expression of all maintained Kazald genes of a species in different parts of their brains, and eye tissue when available. *Kazald1*, *Kazald2*, and *Kazald3* were duplicated in the lineage leading to paddlefish with no subsequent losses, creating a and b versions of these genes. *Kazald2* was duplicated in the lineage leading to eel with no subsequent loss, creating a *Kazald2a* and *Kazald2b*. PRJ IDs indicate the publicly available RNA-Seq datasets which generated the raw data for the listed tissues. Dots represent biological replicates in examined datasets, error bars represent standard deviation when calculable. CPM = Counts Per Million.

**Supplementary Figure 6.**
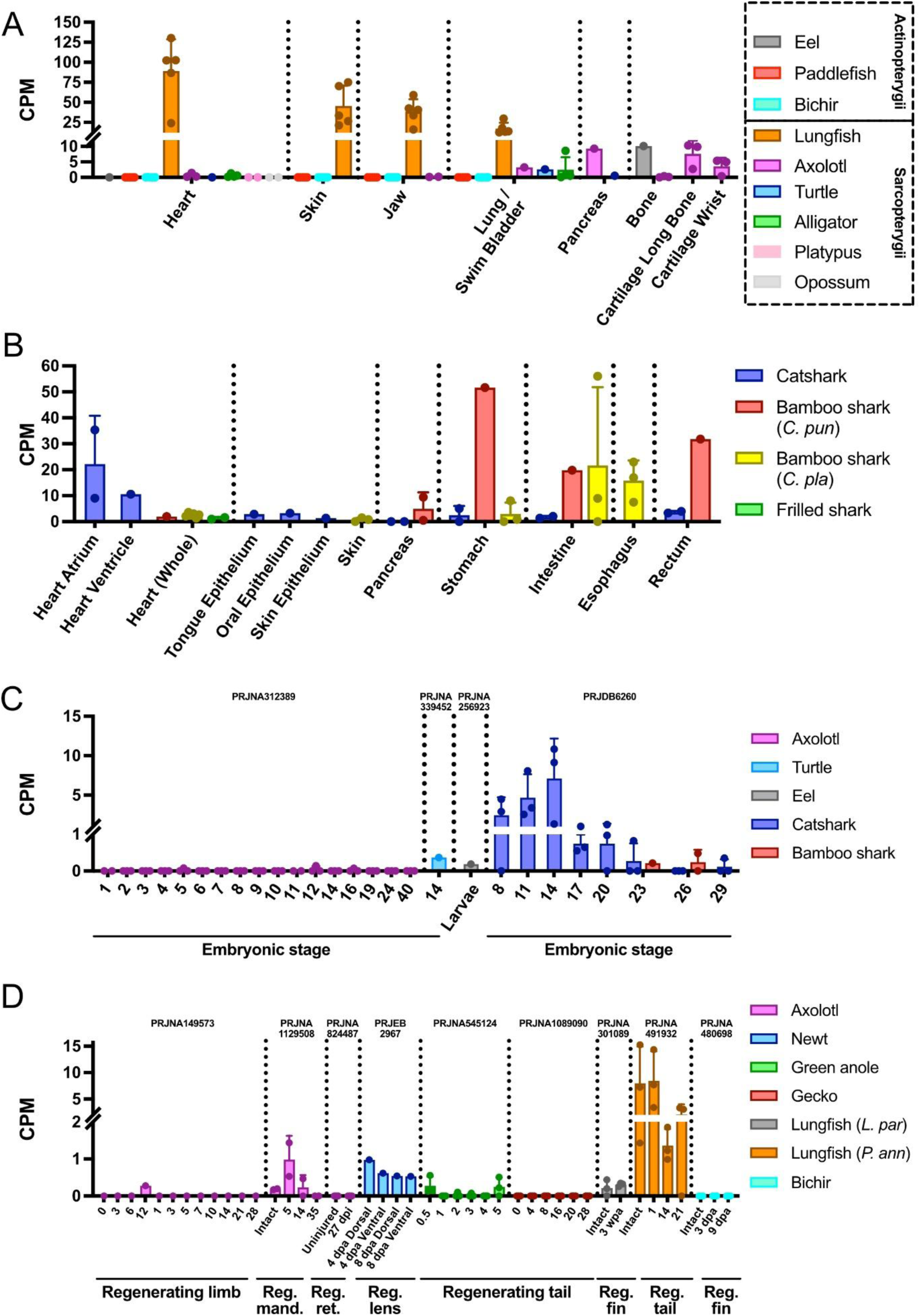
*Kazald4* lacks a conserved expression pattern across jawed vertebrates. A) Quantification of *Kazald4* expression in different tissues of multiple sarcopterygian and actinopterygian species. Data is from PRJNA256923, PRJNA599026, PRJNA300706, PRJNA339452, PRJNA556093, PRJNA143627, PRJNA1129508, and PRJNA378982. B) Quantification of *Kazald4* expression in different tissues of multiple chondrichthyan species. Data is from PRJDB6260, PRJNA1026724, and PRJDB14248. C) Quantification of *Kazald4* expression in whole embryos during embryonic development of multiple species of jawed vertebrates. D) Quantification of *Kazald4* expression during regeneration in different tissues of multiple species of bony vertebrates. PRJ IDs indicate the publicly available RNA-Seq datasets which generated the raw data for the listed tissues. Dots represent biological replicates in examined datasets, error bars represent standard deviation when calculable. CPM = Counts Per Million, Reg. = regenerating, mand. = mandible, ret. = retina.

**Supplementary Figure 7.**
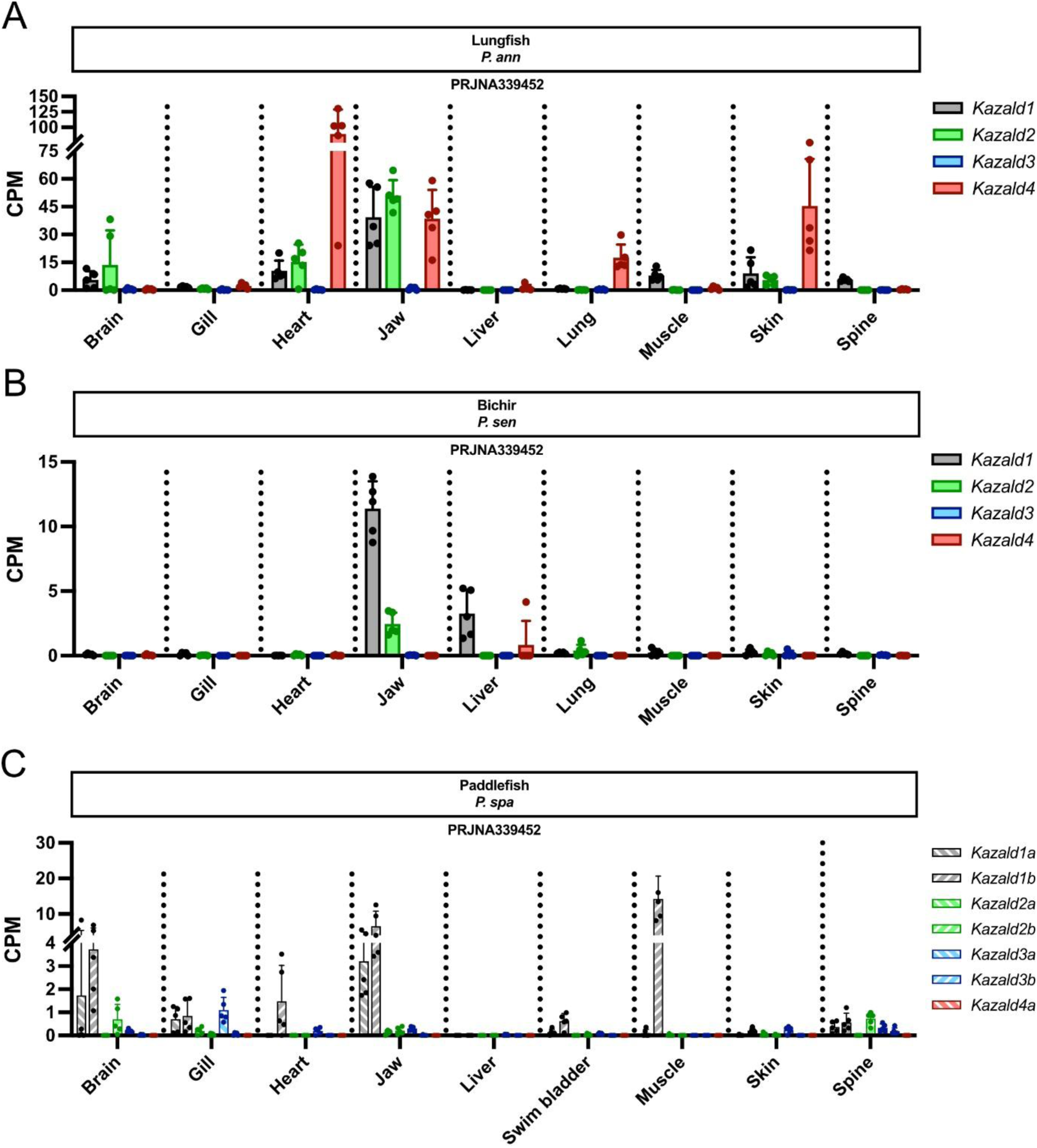
Kazald gene expression in various tissues of non-tetrapod/teleost fish. A) Quantification of the expression of all Kazald genes in various tissues of the lungfish. B) Quantification of the expression of all Kazald genes in various tissues of the bichir. C) Quantification of the expression of all Kazald genes in various tissues of the paddlefish. PRJ IDs indicate the publicly available RNA-Seq datasets which generated the raw data for the listed tissues. Dots represent biological replicates in examined datasets, error bars represent standard deviation when calculable. CPM = Counts Per Million.

**Supplementary Figure 8.**
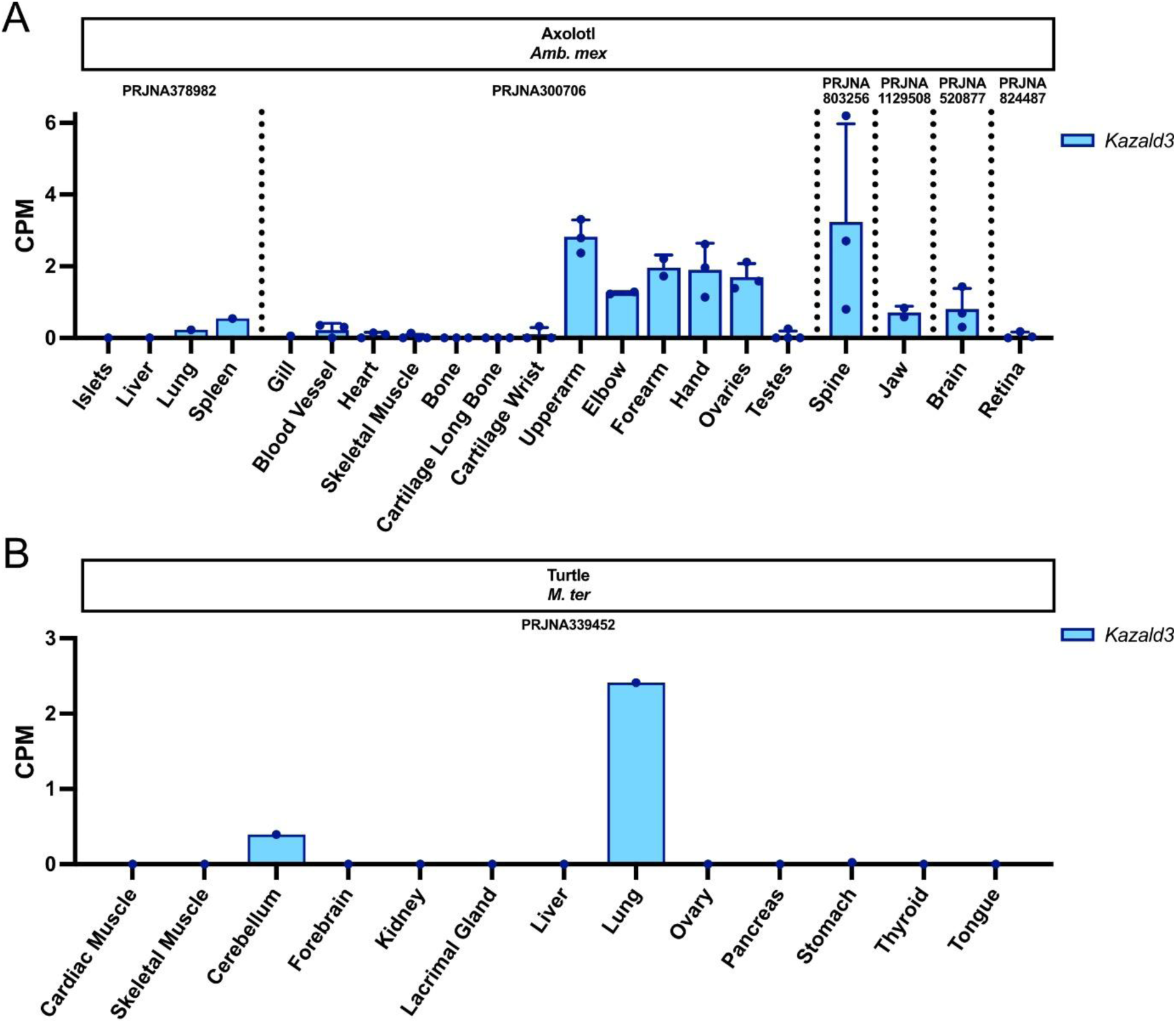
*Kazald3* has no to very low levels of expression in the tissues of adult tetrapods. A) Quantification of *Kazald3* expression in various tissues of the axolotl. B) Quantification of *Kazald3* expression in various tissues of the turtle. PRJ IDs indicate the publicly available RNA-Seq datasets which generated the raw data for the listed tissues. Dots represent biological replicates in examined datasets, error bars represent standard deviation when calculable. CPM = Counts Per Million.

**Supplementary Table 1.** TBLASTN search of the axolotl genome reveals four regions spread across four chromosomes with high similarity to known Kazald genes. Scores are given for the individual exons of the Kazald genes. Identities are number of identical amino acids. Positives are aligned amino acids that are either identical or have similar chemical properties. E Value is the number of expected hits of similar quality that could be found just by chance. Exon 4 is not included due to failure to find a match in the genome.

| **Mouse *Kazald1*** |  |  |  |  |  |  |  |
| --- | --- | --- | --- | --- | --- | --- | --- |
| Axolotl Chromosome | Exon | Start | End | Direction | Identities | Positives | E Value |
| Chromosome 8 | Exon 1 | 1,479,087,933 | 1,479,088,340 | - | 91/137 (66%) | 108/137 (79%) | 7e-54 |
|  | Exon 2 | 1,479,013,148 | 1,479,013,306 | - | 48/53 (91%) | 50/53 (94%) | 4e-26 |
|  | Exon 3 | 1,478,969,760 | 1,478,969,906 | - | 35/49 (71%) | 43/49 (88%) | 7e-19 |
| Chromosome 3q | Exon 1 | 561,620,191 | 561,620,586 | + | 74/135 (55%) | 94/135 (70%) | 2e-41 |
|  | Exon 2 | 561,884,219 | 561,884,377 | + | 31/53 (58%) | 36/53 (68%) | 2e-13 |
|  | Exon 3 | 561,933,219 | 561,933,356 | + | 27/46 (59%) | 31/46 (67%) | 5e-12 |
| Chromosome 10 | Exon 1 | 222,256,888 | 222,257,250 | - | 63/123 (51%) | 81/123 (66%) | 3e-26 |
|  | Exon 2 | 222,122,948 | 222,123,106 | - | 32/53 (60%) | 40/53 (75%) | 1e-15 |
|  | Exon 3 | 222,098,653 | 222,098,790 | - | 23/46 (50%) | 31/46 (67%) | 3e-09 |
| Chromosome 6q | Exon 1 | 1,617,595,903 | 1,617,596,295 | - | 63/132 (48%) | 82/132 (62%) | 2e-27 |
|  | Exon 2 | 1,617,577,351 | 1,617,577,509 | - | 25/53 (47%) | 37/53 (70%) | 2e-10 |
|  | Exon 3 | 1,617,560,918 | 1,617,561,055 | - | 24/46 (52%) | 32/46 (70%) | 8e-11 |
| **Zebrafish *kazald2*** |  |  |  |  |  |  |  |
| Axolotl Chromosome | Exon | Start | End | Direction | Identities | Positives | E Value |
| Chromosome 6q | Exon 1 | 1,617,595,903 | 1,617,596,289 | - | 68/136 (50%) | 80/136 (59%) | 9e-28 |
|  | Exon 2 | 1,617,577,351 | 1,617,577,509 | - | 25/53 (47%) | 37/53 (70%) | 1e-10 |
|  | Exon 3 | 1,617,560,921 | 1,617,561,058 | - | 27/46 (59%) | 36/46 (78%) | 4e-13 |
| Chromosome 3q | Exon 1 | 561,620,212 | 561,620,583 | + | 57/132 (43%) | 77/132 (58%) | 8e-23 |
|  | Exon 2 | 561,884,219 | 561,884,377 | + | 23/53 (43%) | 35/53 (66%) | 1e-10 |
|  | Exon 3 | 561,933,216 | 561,933,353 | + | 27/46 (59%) | 35/46 (76%) | 1e-13 |
| Chromosome 8 | Exon 1 | 1,479,087,933 | 1,479,088,313 | - | 55/134 (41%) | 77/134 (57%) | 3e-21 |
|  | Exon 2 | 1,479,013,148 | 1,479,013,306 | - | 25/53 (47%) | 32/53 (60%) | 3e-09 |
|  | Exon 3 | 1,478,969,769 | 1,478,969,903 | - | 26/45 (58%) | 34/45 (76%) | 2e-11 |
| Chromosome 10 | Exon 1 | 222,256,888 | 222,257,241 | - | 60/123 (49%) | 73/123 (59%) | 2e-19 |
|  | Exon 2 | 222,122,948 | 222,123,106 | - | 22/53 (42%) | 31/53 (58%) | 3e-09 |
|  | Exon 3 | 222,098,656 | 222,098,793 | - | 25/46 (54%) | 34/46 (74%) | 6e-11 |

**Supplementary Table 2.** Most jawed vertebrates possess multiple Kazald genes. Putative Kazald genes from the species which possessed them were used for the creation of the large-scale phylogenetic tree (Fig. 1). Bolded letters in the column “Scientific Name” are used as the IDs of species in the tree. The column “Number of Manually Found or Edited Genes” refers to genes either not in the transcriptome, or manually edited based on the conserved Kazald gene genomic profile that we identified (see Materials and Methods). Use of – indicates absence.

| Common Name | Scientific Name | Total Number of Kazald Genes | Number of Manually Found or Edited Genes |
| --- | --- | --- | --- |
| Sponge | *E. muelleri* | – | – |
| Sponge | *O. lobularis* | – | – |
| Placozoa | *T. adhaerens* | – | – |
| Hydra | *H. vulgaris* | – | – |
| Staghorn Coral | *A. muricata* | – | – |
| Starlet Sea Anemone | *N. vectensis* | – | – |
| Flame Jellyfish | *R. esculentum* | – | – |
| Fruit Fly | *D. melanogaster* | – | – |
| Nematode | *C. elegans* | – | – |
| Planaria | *S. mediterranea* | – | – |
| Planaria | *D. japonica* | – | – |
| Milk-White Planarian | *D. lacteum* | 1<sup>*,**</sup> | – |
| Atlantic Horseshoe Crab | ***L. polyphemus*** | 1<sup>***</sup> (3) | – |
| Cactus Worm | ***P. caudatus*** | 1 | – |
| Portuguese Oyster | ***C. angulata*** | 1 | – |
| Leech | ***H. robusta*** | 1 | – |
| Acorn Worm | ***S. kowalevskii*** | 1 | – |
| Crown-of-Thorns Starfish | ***A. planci*** | 1 | – |
| Florida Lancelet | ***B. floridae*** | 1 | – |
| Inshore Hagfish | *E. burgeri* | 1 | – |
| Sea Lamprey | *P. marinus* | 1 | – |
| Elephant Shark | *C. milii* | 2 | – |
| Great White Shark | *C. carcharias* | 2 | – |
| Thorny Skate | *A. radiata* | 2 | – |
| Gray Bichir | *P. senegalus* | 4 | – |
| Sterlet | *A. ruthenus* | 7 | 2 |
| American Paddlefish | *P. spathula* | 7 | 2 |
| Spotted Gar | *L. oculatus* | 3 | – |
| European Eel | *A. anguilla* | 4 | 1 |
| Atlantic Tarpon | *M. atlanticus* | 4 | 3 |
| Asian Bonytongue | *S. formosus* | 2 | – |
| Atlantic Herring | *C. harengus* | 3 | 1 |
| Allis Shad | *A. alosa* | 3 | 1 |
| Mexican Tetra | *Ast. mexicanus* | 3 | 1 |
| Red-Bellied Piranha | *P. nattereri* | 3 | – |
| Zebrafish | *D. rerio* | 2 | – |
| Northern Pike | *E. lucius* | 2 | 1 |
| Chinook Salmon | *O. tshawytscha* | 2 | – |
| Atlantic Cod | *G. morhua* | 2 | – |
| European Seabass | *D. labrax* | 2 | – |
| Torafugu Pufferfish | *T. rubripes* | 1 | – |
| Medaka | *O. latipes* | 1 | – |
| Coelacanth | *L. chalumnae* | 3 | – |
| West African Lungfish | *P. annectens* | 4 | – |
| Axolotl | *Amb. mexicanum* | 4 | – |
| Iberian Ribbed Newt | *P. waltl* | 3**** (4) | – |
| Eastern Newt | *N. viridescens* | 3** | – |
| Ramos' Mushroom-Tongue Salamander | *B. ramosi* | 4** | – |
| Two-Lined Caecilian | *R. bivittatum* | 3 | 2 |
| Gaboon Caecilian | *G. seraphini* | 2 | 1 |
| Tiny Cayenne Caecilian | *M. unicolor* | 2 | – |
| European Fire-Bellied Toad | *B. bombina* | 3 | – |
| Western Clawed Frog | *X. tropicalis* | 1 | – |
| Common Toad | *B. bufo* | 2 | – |
| Common Frog | *R. temporaria* | 2 | – |
| Painted Turtle | *C. picta* | 3 | – |
| Goode's Thornscrub Tortoise | *G. evgoodei* | 3 | – |
| Green Sea Turtle | *C. mydas* | 2 | – |
| Yangtze Giant Softshell Turtle | *R. swinhoei* | 2 | 2 |
| Red-Bellied Short-Necked Turtle | *E. subglobosa* | 2 | 2 |
| Gharial | *G. gangeticus* | 2 | – |
| Chinese Alligator | *A. sinensis* | 2 | – |
| Emu | *Dro. novaehollandiae* | 2 | – |
| Chicken | *G. gallus* | 2 | – |
| Zebra Finch | *T. guttata* | 2 | – |
| Tuatara | *S. punctatus* | 3 | 2 |
| Leopard Gecko | *E. macularius* | 2 | – |
| Sand Lizard | *L. agilis* | 2 | – |
| Eastern Fence Lizard | *S. undulatus* | 2 | – |
| Western Terrestrial Garter Snake | *T. elegans* | 1 | – |
| Platypus | *O. anatinus* | 2 | – |
| Short-Beaked Echidna | *T. aculeatus* | 2 | 1 |
| Gray Short-Tailed Opossum | *M. domestica* | 2 | 1 |
| Common Brushtail Possum | *T. vulpecula* | 2 | – |
| Tasmanian Devil | *S. harrisii* | 2 | 1 |
| Indian Elephant | *E. max. indicus* | 1 | – |
| Nine-Banded Armadillo | *Das. novemcinctus* | 1 | – |
| Horse | *E. caballus* | 1 | – |
| Large Flying Fox | *P. vampyrus* | 1 | – |
| Dog | *C. lup. familiaris* | 1 | – |
| Mouse | *M. musculus* | 1 | – |
| Human | *H. sapiens* | 1 | – |

\* = not used in the large-scale phylogenetic tree
\*\* = only transcriptomes are available, thus the total number of Kazald genes may be greater
\*\*\* = 3 potential Kazald genes found broken up across scaffolds, thus only the 1 fully intact gene used in the phylogenetic tree
\*\*\*\* = 4 Kazald genes found in the genome, but only the 3 genes in the transcriptome used in the phylogenetic tree

